# Identification and Characterization of Fbxl22, a novel skeletal muscle atrophy-promoting E3 ubiquitin ligase

**DOI:** 10.1101/2020.04.24.059659

**Authors:** David C. Hughes, Leslie M. Baehr, Julia R. Driscoll, Sarah A. Lynch, David S. Waddell, Sue C. Bodine

## Abstract

Muscle-specific E3 ubiquitin ligases have been identified in muscle atrophy-inducing conditions. The purpose of the current study was to explore the functional role of Fbxl22, and a newly identified splice variant (Fbxl22-193), in skeletal muscle homeostasis and neurogenic muscle atrophy. In mouse C_2_C_12_ muscle cells, promoter fragments of the Fbxl22 gene were cloned and fused with the secreted alkaline phosphatase reporter gene to assess the transcriptional regulation of Fbxl22. The tibialis anterior muscles of male C57/BL6 mice (12-16 weeks old) were electroporated with expression plasmids containing the cDNA of two Fbxl22 splice variants and tissues collected after 7, 14 and 28 days. Gastrocnemius muscles of wild type and MuRF1 knockout mice were electroporated with an Fbxl22 RNAi or empty plasmid, denervated three days post-transfection, and tissues collected 7 days post-denervation. The full-length gene and novel splice variant are transcriptionally induced early (after 3 days) during neurogenic muscle atrophy. In vivo overexpression of Fbxl22 isoforms in mouse skeletal muscle lead to evidence of myopathy/atrophy suggesting that both are involved in the process of neurogenic muscle atrophy. Knockdown of Fbxl22 in MuRF1 KO muscles resulted in significant additive muscle sparing at 7 days of denervation. Targeting two E3 ubiquitin ligases appears to have a strong additive effect on protecting muscle mass loss with denervation and these findings have important implications in the development of therapeutic strategies to treat muscle atrophy.

## Introduction

Skeletal muscle wasting can occur as a result of both internal (e.g. genetic disorders) and external stimuli (e.g. disuse) (16, 23, 69). Central to the process of muscle wasting/atrophy is increased protein degradation through the ubiquitin-proteasome system (UPS) and autophagosome lysosome system (5, 8, 55). The UPS targets proteins for degradation by the 26S proteasome. The ubiquitin ligase enzyme, or E3, is critical for catalyzing the movement of ubiquitin from the E2 to the protein substrate which can lead to altered function or degradation by the proteasome (5, 55). There are approximately 40 E2 enzymes and over 600 E3 ubiquitin ligases in the human genome making it an extremely complex and relatively understudied system (5, 35). In skeletal muscle, only a few E3 ubiquitin ligases have been characterized and associated with muscle atrophy, including MuRF1 (muscle-specific RING finger 1), MAFbx (muscle atrophy F-box; also known as Atrogin-1) (9, 28), MUSA1 (muscle ubiquitin ligase of the SCF complex in atrophy-1) (57) and SMART (Specific of Muscle Atrophy and Regulated by Transcription) (45). Recently, we identified UBR5 as an E3 ubiquitin ligase that may play a role in muscle growth and hypertrophy (59). Considerably more E3 ubiquitin ligases are expressed in skeletal muscle, most of which remain to be explored in the context of muscle physiology and disease.

Of all the E3 ubiquitin ligases, MuRF1 and MAFbx have been the most studied, especially during the progression of skeletal muscle atrophy induced by a wide range of stressors (such as neural inactivity, starvation, and aging) since their discovery in 2001 (9, 28). Under certain atrophy stimuli, increased MuRF1 and MAFbx expression occurs via activation of the glucocorticoid receptor and/or FoxO transcription factors (56, 66). A null deletion in MuRF1 and MAFbx leads to significant muscle sparing after 14 days of neural inactivity (9). Interestingly, after 7 days of neural inactivity, MuRF1 knockout (KO) animals display muscle atrophy similar to wild type animals, suggesting other unidentified factors contribute to the process of skeletal muscle wasting (9, 21). In our previously published microarray data set, 400-600 genes showed significant up-regulation in wild type and/or MuRF1 KO skeletal muscle exposed to neural inactivity for 3 days (25). Of the 400-600 genes identified, many were originally identified by the RIKEN project and thus of unknown name and function at the time of the study (36).

One such RIKEN gene from the microarray data set was subsequently identified to be an E3 ubiquitin ligase from the F-box family, named F-box and leucine-rich protein 22 (Fbxl22) (25). In cardiac muscle, Fbxl22 is enriched and, similar to MAFbx, a component of the Skp1-Cullin-F-box (SCF) ubiquitin ligase complex through its N-terminal F-box domain (9, 28, 62). Using a yeast 2-hybrid screen and cardiomyocytes, Fbxl22 has been suggested to target filamin-C and α-actinin-2 for proteasomal degradation (62). Filamin-C and α-actinin-2 are sarcomeric proteins localized to the Z-line and are critical for maintaining the spatial relationship between myofilaments and stabilizing the myofilament lattice (4, 34, 39). Little is known about the role of Fbxl22 in skeletal muscle and its involvement in the atrophy process. Therefore, we sought to characterize Fbxl22 expression and function in muscle cells and investigate the role of Fbxl22 in skeletal muscle homeostasis and neural inactivity-induced atrophy.

## Materials and methods

### Cell Culture

Cell culture was performed as previously described (29, 30). Briefly, mouse C_2_C_12_ myoblasts were obtained from the American Type Culture Collection (Manassas, VA) and maintained in Dulbecco’s Modified Eagle’s Medium (DMEM) (Thermo Fischer Scientific, Waltham, MA) supplemented with 10% Fetal Bovine Serum (FBS) (GE Healthcare HyClone Laboratories, Logan, UT), non-essential amino acids (GE Healthcare HyClone Laboratories, Logan, UT), 1X Penicillin/Streptomycin and Gentamicin (Thermo Fischer Scientific, Waltham, MA). C_2_C_12_ cells were maintained in a humidified chamber at 37°C and 5% CO_2_ and grown to 100% confluency and then switched to DMEM supplemented with 2% serum to induce myogenic differentiation.

### Secreted Alkaline Phosphatase (SEAP) reporter gene assays

SEAP reporter assays were performed as previously described (29, 30, 42). All reporter assays were conducted in C_2_C_12_ cells cultured in 12-well plates at a seeding density of 100,000 cells/well. Cells were transfected at 70-80% confluency with 1 μg of total DNA/well (250 ng reporter plasmid, 125 ng internal control plasmid, 125–250 ng of expression plasmids, and empty pBluescript vector as filler DNA to 1 μg/well). DNA was complexed with Turbofect transfection reagent (Thermo Fisher Scientific, Waltham, MA) for 20 minutes prior to overlaying of cells according to the manufacturer’s protocol. The media was sampled at 24-hours post-transfection and then switched to DMEM supplemented with 2% serum to induce differentiation. Media was sampled at 2- and 7-days post-media change and reporter activity was assessed using the 3-(1-chloro-3′-methoxyspiro-(adamantane-4,4′-dioxetane)-3′-yl) phenyl dihydrogen phosphate (CSPD) chemiluminescent substrate and the Phospha-Light SEAP Reporter Gene Assay System according to the manufacturer’s protocol (Thermo Fisher Scientific, Waltham, MA). Luminescence levels were detected using a Synergy 2 microplate reader set for an endpoint read with a 2s integration time. At the end of the time course, cells were lysed with Passive Lysis Buffer (Promega, Madison, WI) and homogenates were cleared by centrifugation and analyzed for β-galactosidase activity. β-galactosidase activity was determined by incubating cell lysates with ortho-nitrophenyl-β-D-galactopyranoside (ONPG) dissolved in Z Buffer (60 mM Na2HPO4·7H2O, 40 mM NaH2PO4·H2O, 10 mM KCl, 1 mM MgSO4, 50 mM β-mercaptoethanol, pH 7.0) overnight at 37°C and subsequently read using a Synergy 2 plate reader at 420nm. Reporter activity was expressed as relative light units of SEAP divided by β-galactosidase activity.

### RNA isolation and cDNA synthesis from C_2_C_12_ cells

Total RNA was isolated from C_2_C_12_ myoblasts and differentiated myotubes using the RNeasy Mini kit (Qiagen, Valencia, CA) according the manufacturer’s protocol. Purified total RNA was reverse transcribed to cDNA using M-MLV Reverse Transcriptase and a Poly T primer according to the manufacturer’s instructions (Thermo Fisher Scientific, Waltham, MA).

### Cloning of a novel Fbxl22 cDNA and promoter fragments

A novel open reading frame of the Fbxl22 gene (GenBank Accession Number: MT011389), termed Fbxl22-193 in this study, was amplified using cDNA generated from RNA isolated from C_2_C_12_ cells as described above (Thermo Fisher Scientific, Waltham, MA). Using the cDNA primers listed in Table 1, the Fbxl22-193 cDNA was amplified and subcloned into the HindIII and EcoRI sites of pcDNA3.1(+) (ThermoFisher Scientific, Waltham, MA). A pCMV6-myc-DDK-Fbxl22 full-length cDNA (GenBank Accession Number: NM_175206) plasmid was purchased from Origene (Rockville, MD). The full-length Fbxl22 variant was termed Fbxl22-236 in this study. An untagged version of the full-length mouse Fbxl22 gene was generated by cutting the full-length Fbxl22-myc-DDK cDNA out of pCMV6 and cloning it into the EcoRI and NotI sites of pcDNA3.1 and then performing site-directed mutagenesis to introduce a stop codon immediately upstream of the myc and DDK tags. The untagged full-length Fbxl22 cDNA was then sequenced to confirm correct orientation, the successful introduction of the correct stop codon, and the absence of any other mutations (Eurofins Genomics, Louisville, KY).

The Fbxl22 proximal promoter region was amplified from genomic DNA isolated from C_2_C_12_ cells using the DNeasy Blood and Tissue Kit (Qiagen, Valencia, CA) according to the manufacturer’s protocol. Fbxl22 promoter specific primers were designed using genomic DNA sequences on deposit with the Ensembl database to amplify approximately 500 and 1000bp fragments of the proximal 5’ regulatory region (Table 1). Promoter fragments were amplified using 1 μg of genomic DNA and Taq Polymerase according to the manufacturer’s protocol (Thermo Fisher Scientific, Waltham, MA). Amplified fragments were subcloned into the MluI and EcoRI sites of the pSEAP2-Basic reporter plasmid (Clontech, Mountain View, CA) and sequenced to confirm correct orientation (Eurofins Genomics, Louisville, KY).

### Site-directed mutagenesis

Site-directed mutagenesis was used to mutagenize conserved putative Ebox sequences in the Fbxl22 proximal promoter and to introduce a stop codon in the pCMV6-myc-DDK-Fbxl22 plasmid using the primers listed in Table 1. The QuikChange II Site-Directed Mutagenesis protocol was performed using 200 ng of plasmid DNA and PfuTurbo DNA Polymerase according to the manufacturer’s protocol (Agilent Technologies, Santa Clara, CA). Mutagenesis reactions were then digested with Dpn I (New England Biolabs, Ipswich, MA) and transformed into sub-cloning efficiency competent cells (New England Biolabs, Ipswich, MA). All constructs were sequenced to confirm successful generation of the expected mutation (Eurofins Genomics, Louisville, KY).

### Quantitative RT-PCR analysis of Fbxl22 expression in cell culture

Total RNA was isolated from C_2_C_12_ cells using the RNeasy minikit (Qiagen, Valencia, CA) and reverse transcribed into cDNA using 1 μg of RNA and iScript Reverse Transcriptase (Bio-Rad, Hercules, CA) according to the manufacturer’s protocol. Quantitative polymerase chain reaction was performed with 100 ng of cDNA, 100 nM of each qPCR primer listed in Table 1, and iTaq Universal SYBR Green Supermix (Bio-Rad, Hercules, CA) according to the manufacturer’s protocol. Relative Fbxl22 expression levels were determined using a Bio-Rad CFX Connect machine and normalized to GAPDH expression using the −ΔΔC_t_ method.

### Study approval for animal experiments

The mice used in these studies were males from the C57BL/6 strain obtained from Charles River laboratories at ages 3-4 months and used for experiments within 2 weeks of their arrival. Male MuRF1 knockout (KO) mice were obtained from a breeding colony maintained at the University of Iowa at 6 months of age and used for denervation studies. The generation of mice with a null deletion of MuRF1 has been previously described (9). Animals were housed in ventilated cages maintained in a room at 21 °C with 12-h light/ dark cycles and had *ad libitum* access to standard chow (Harlan-Teklad formula 7913) and water throughout the study. All animal procedures were approved by the Institutional Animal Care and Use Committee at the University of Iowa.

### Denervation Model

Targeted denervation of the lower limb muscles in the right leg was accomplished through transection of the sciatic nerve as previously described (27). Under isoflurane anesthesia (2-4% inhalation) and with the use of aseptic surgical techniques, the sciatic nerve was isolated in the midthigh region and cut with sharp scissors. Mice were given an analgesic (buprenorphine, 0.1 mg/kg) immediately following the surgery and returned to their cage following recovery. A group of true control animals were used for the in-depth time course analysis of muscle mass plus MuRF1 and Fbxl22 isoform mRNA expression.

### In vivo electroporation

Transfection of mouse skeletal muscle with plasmid DNA was performed in mice under isoflurane (2-4% inhalation) anesthesia as previously described (22, 59). Briefly, after a 2hr pre-treatment with 0.4 units/μl of bovine placental hyaluronidase (Sigma) resuspended in sterile 0.9% saline, 20μg of plasmid DNA was injected into either the tibialis anterior (TA) or the lateral and medial gastrocnemius muscles (lateral gastrocnemius, LGA; medial gastrocnemius, MGA). The hind limbs were placed between two-paddle electrodes and subjected to 10 pulses (20 msec) of 175 V/cm using an ECM-830 electroporator (BTX Harvard Apparatus, Holliston, MA). For in vivo overexpression studies, an additional 2 μg of emerald-GFP plasmid was electroporated for identification of GFP positive fibers. For RNAi plus denervation experiments, plasmids were electroporated into hindlimb muscles three days prior to implementation of sciatic nerve surgery (detailed above). Following transfection, mice were returned to their cages to resume normal activities until tissue collection.

### Tissue Collection

Following completion of the appropriate time period, mice were anesthetized with isoflurane, and the TA and GA muscles were excised, weighed, frozen in liquid nitrogen, and stored at −80°C for later analysis. A subset of muscles were collected for histology and processed as described below. On completion of tissue removal, mice were euthanized by exsanguination.

### Histology

Harvested muscles were immediately fixed in 4% (w/v) paraformaldehyde for 16h at 4°C. Following a sucrose gradient incubation period, the TA muscles were embedded in Tissue Freezing Medium (Triangle Biomedical Sciences), and a Thermo HM525 cryostat was used to prepare 10-μm serial sections from the muscle midbelly. All sections were examined and photographed using a Nikon Eclipse Ti automated inverted microscope equipped with NIS-Elements BR digital imaging software.

#### Laminin Stain

TA muscle sections were permeabilized in PBS with 1% triton for 10 minutes at room temperature. After washing with PBS, sections were blocked with 5% goat serum for 15 minutes at room temperature. Sections were incubated with Anti-Laminin (1:500, Sigma Aldrich, Cat no. L9393) in 5% goat serum for 2 hours at room temperature, followed by two 5-minute washes with PBS. Goat-anti-rabbit AlexaFluor® 555 secondary (1:333) in 5% goat serum was then added for 1 hour at room temperature. Slides were cover slipped using ProLong Gold Antifade reagent (Life Technologies). Image analysis was performed using Myovision software (68). Skeletal muscle fiber size was analyzed by measuring ≥ 350 transfected muscle fibers per muscle (GFP-positive), per animal (10x magnification). In Fbxl22 overexpression studies, we also measured the size of non-transfected fibers (GFP-negative) from the transfected muscles at all time points. Comparison of the CSA distributions of GFP-negative fibers in EV, −236 and 193 transfected muscles revealed no difference between groups. Further in some muscles the transfection frequency was so high that there were few GFP-negative fibers within the same region as the GFP-positive fibers. Therefore, for Fbxl22 overexpression studies, fiber size comparisons were made between the GFP-positive fibers in the EV controls and the −293 and 193 transected muscles.

#### Hematoxylin & Eosin stain

A standard H&E protocol was performed (adapted from Wang et al. (67)). Briefly, TA muscle sections were incubated with hematoxylin solution for 10 minutes to stain the nuclei and then stained with Eosin solution for 3 min. Sections were then immersed in 70% ethanol for 20 sec, 90% ethanol for 20 sec, 100% ethanol for 1 min, and citrisolv for 3 min. Finally, slides were allowed to air dry and a coverslip was added using Permount media (Thermo Fisher Scientific, Waltham, MA). Percentage of centrally nucleated myofibers was calculated by dividing the number of transfected fibers containing ≥1 centrally located nucleus by the total number of transfected fibers (as previously described (24)). An average of 1500 myofibers were counted per animal (N=4-5/group), and quantification was performed using ImageJ software (National Institutes of Health, Bethesda, MD, USA).

### RNAi Plasmids

For RNAi experiments, the negative control RNAi plasmid was described previously (22, 59) and encodes emerald green fluorescent protein (EmGFP) and a non-targeting pre-miRNA under bicistronic control of the cytomegalovirus (CMV) promoter in the pcDNA6.2GW/EmGFP-miR plasmid (Invitrogen, Carlsbad, CA). The Fbxl22 RNAi plasmid encodes EmGFP and an artificial pre-miRNA targeting the full length gene of mouse Fbxl22 under bicistronic control of the CMV promoter; it was generated by ligating the Mmi576126 oligonucleotide duplex (Invitrogen, Carlsbad, CA) into the pcDNA6.2GW/EmGFP-miR plasmid.

### Immunoblotting

Frozen TA and GA muscles were homogenized in sucrose lysis buffer (50 mM Tris pH 7.5, 250 mM sucrose, 1 mM EDTA, 1 mM EGTA, 1% Triton X 100, 50 mM NaF). The supernatant was collected following centrifugation at 8,000 *g* for 10 minutes and protein concentrations were determined using the 660 protein assay (Thermo Fisher Scientific, Waltham, MA). Twelve micrograms of protein were subjected to SDS-PAGE on 4-20% Criterion TGX stain-free gels (Bio-Rad, Hercules, CA) and transferred to polyvinylidene diflouride membranes (PVDF, Millipore, Burlington, MA). Membranes were blocked in 3% nonfat milk in Tris-buffered saline with 0.1% Tween-20 added for one hour and then probed with primary antibody overnight at 4°C. Membranes were washed and incubated with HRP-conjugated secondary antibodies at 1:10,000 for one hour at room temperature. Immobilon Western Chemiluminescent HRP substrate was then applied to the membranes prior to image acquisition. Image acquisition and band quantification were performed using the Azure C400 System (Azure Biosystems, Dublin, CA, USA) and Image Lab, version 6.0.1 (Bio-Rad), respectively. Total protein loading of the membranes captured from images using stain-free gel technology was used as the normalization control for all blots. The following antibodies were used: Ubiquitin (1:1000, Life Sensors, VU-1), p62 (1:1000, Sigma, P0067), LC3B (1:1000, Sigma, L7543), Desmin (1:500, Cell Signaling, 5332), Dystrophin (1:1000, Santa Cruz, sc-365954), α-actinin (1:1000, Cell Signaling, 3134), Vimentin (1:1000, Santa Cruz, sc-373717).

### Gene expression by Quantitative RT -PCR in skeletal muscle

Frozen muscle powder was homogenized using RNAzol RT reagent (Sigma-Aldrich, St Louis, MO) in accordance with the manufacturer’s instructions. cDNA was synthesized using a reverse transcription kit (iScript cDNA synthesis kit; Bio-Rad, Hercules, CA) from 1 μg of total RNA. PCR reactions (10 μL) were set up as: 2 μL of cDNA, 0.5 μL (10 μM stock) forward and reverse primer, 5 μL of Power SYBR Green master mix (Thermo Fisher Scientific, Waltham, MA) and 2 μL of RNA/DNA free water. Gene expression analysis was then performed by quantitative PCR on a Quantstudio 6 Flex Real-time PCR System (Applied Biosystems, Foster City, CA) using the mouse primers shown in Table 1. PCR cycling comprised: hold at 50°C for 5 min, 10 min hold at 95°C, before 40 PCR cycles of 95°C for 15 s followed by 60°C for 1 min (combined annealing and extension). Melt curve analysis at the end of the PCR cycling protocol yielded a single peak. As a result of reference gene instability, gene expression was normalized to tissue weight and subsequently reported as the fold change relative to control muscles, as described previously (31). This type of analysis has previously been used extensively by our group (2, 3, 32).

### Statistical Analysis

Results are presented as mean ± standard deviation (SD) for all in vitro experiments and mean ± standard error of measure (SEM) for all in vivo experiments. Statistical differences for all in vitro reporter assays were determined using a two-sample t-test and performed using the data analysis add-in for Excel. Statistical differences for in vivo overexpression studies were determined using a paired students’ t-test. The denervation time course experiment was assessed for differences in muscle mass and mRNA expression using a one-way analysis of variance (ANOVA) with Dunnett’s multiple comparisons test. For the RNAi plasmid plus denervation experiments, differences were determined using an unpaired students’ t-test. All in vivo statistical analyses were performed using GraphPad Prism (GraphPad Software, Inc., La Jolla, CA). Results were considered significant when P ≤ 0.05.

## Results

### Fbxl22 is induced during neurogenic skeletal muscle atrophy

An Illumina beadchip microarray was previously performed on RNA isolated from triceps surae (TS) muscle of mice undergoing denervation-induced skeletal muscle atrophy and described in Furlow *et al*. (25). Further analysis of the data SubSeries (GEO: GSE44205) submitted to the NCBI Gene Expression Omnibus (GEO) revealed numerous genes that previously have not been described in skeletal muscle, but show differential expression following sciatic nerve transection. One of these previously undescribed genes was originally identified from the microarray by its Riken designation, 1110004B15RIK, and later determined to be Fbxl22, an E3 ubiquitin ligase known to be enriched in cardiac muscle (18). Based on the microarray data, basal expression of Fbxl22 is low in skeletal muscle of wild type (WT) and MuRF1 KO mice, but is significantly elevated after three days of denervation (Fig 1A). Interestingly, Fbxl22 expression returns to baseline expression by 14 days of denervation, in both WT and MuRF1 KO skeletal muscle (Fig. 1B).

**Figure 1.**
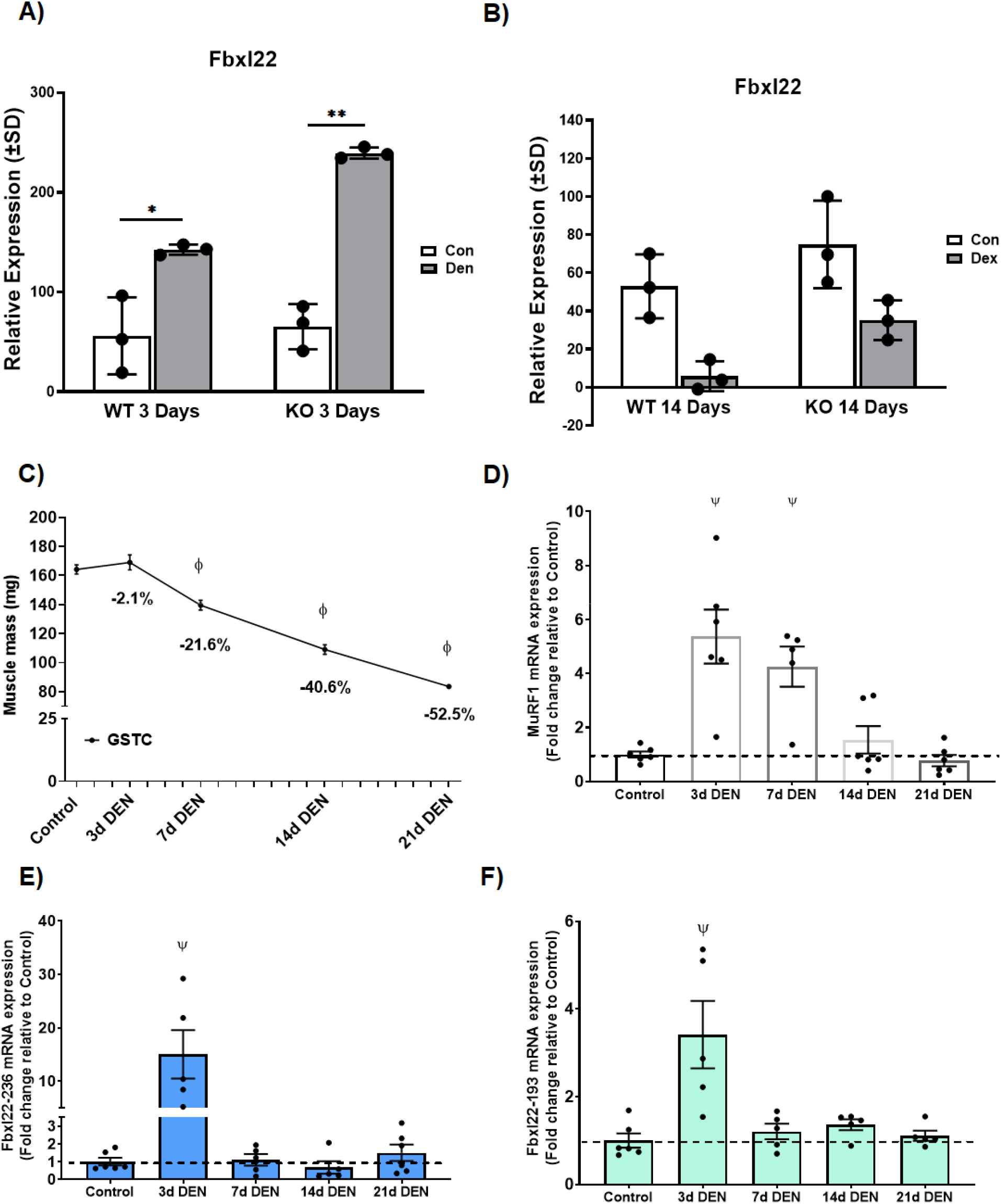
Fbxl22 is induced during neurogenic skeletal muscle atrophy. Whole genome expression analysis was conducted on triceps surae muscle from wild-type (WT) and MuRF1-null (KO) mice after **(A)** 3 days and **(B)** 14 days of denervation (DEN). Fbxl22 expression increased significantly at 3 days of denervation but returned to basal expression levels by 14 days post-denervation. Each condition represents the average expression from three animals (n = 3) and error bars represent ± SD. Gray bars, controls; black bars, DEN. Significant difference between denervated mice and control mice in the same group, (*: *P* ≤ 0.05, **: P ≤ 0.01). Data in Panel A and B from a previously published study (25). (**C**) Muscle mass (milligrams) of Gastrocnemius Complex (GSTC) muscle exposed to denervation (DEN) for 3, 7, 14 and 21 days. The loss of mass as a percentage of control muscles is also given for each time point. (**D**) MuRF1, **(E)** Fbxl22-236 and **(F)** Fbxl22-193 mRNA expression during denervation time course in GSTC muscle. Data presented as Mean ± SEM (N= 5-6/group). ϕ and Ψ depict P ≤ 0.05 verses control group.

To confirm the identification of Fbxl22 from our previously published microarray data (25) and to examine levels of endogenous expression for a novel Fbxl22 splice variant (identification detailed below), we performed an in depth time course of denervation-induced muscle atrophy. For the gastrocnemius complex (lateral and medial gastrocnemius, plantaris, and soleus muscles), we observed significant reductions in muscle mass at 7, 14 and 21 days post sciatic nerve transection (Fig. 1C). A significant increase in MuRF1 mRNA expression occurred at 3 and 7 days post-denervation (Fig. 1D). Interestingly, Fbxl22 (both full length, Fbxl22-236 and novel splice variant, Fbxl22-193) mRNA expression was significantly increased at 3 days post-denervation, but was back to baseline levels by 7 days (Fig. 1E and 1F).

### Identification and cloning of a novel Fbxl22 splice variant from skeletal muscle cells

Fbxl22 is located on chromosome 9 in mouse and encodes two alternative transcripts that consist of either 2 coding exons (Fbxl22-Full Length) or 3 coding exons (Fbxl22-Novel Transcript) (Fig. 2A). To demonstrate that this gene is expressed endogenously at the transcriptional level in muscle cells, a cDNA comprising the open reading frame (ORF) of Fbxl22 was amplified from C_2_C_12_ myoblasts. The amplified cDNA was sequenced in both directions and found to contain an in frame 129 nucleotide deletion in the second exon based on comparisons to the predicted sequence available from the Ensembl database (Fig. 2A). This deletion results in the removal of 43 amino acids in the leucine-rich repeat region of Fbxl22 and essentially removes one leucine-rich repeat. To further assess the endogenous Fbxl22 expression profile in muscle cells, real-time RT-qPCR was performed using RNA isolated from proliferating myoblasts and myotubes at early and late stages of differentiation. The results demonstrate that Fbxl22-193 (novel) and Fbxl22-236 (full length) expression levels are highest in fully differentiated myotubes (DD7) compared to proliferating myoblasts (PD2) (Fig. 2B and 2C). To determine if the Fbxl22-193 splice variant can be generated from the full-length Fbxl22 transcript, C_2_C_12_ cells were transfected with an expression plasmid containing the Fbxl22-236 cDNA and quantitative RT-qPCR was performed using primers specific for Fbxl22-236 or Fbxl22-193. The results show that overexpression of Fbxl22-236 in muscle cells leads to significantly elevated levels of both the full-length Fbxl22 transcript and the novel Fbxl22-193 splice variant (Fig. 2D and 2E). Furthermore, this observation was confirmed in vivo with Fbxl22-236 overexpression in mouse skeletal TA muscle leading to significantly increased levels of the novel Fbxl22-193 splice variant (Figure S1).

**Figure 2.**
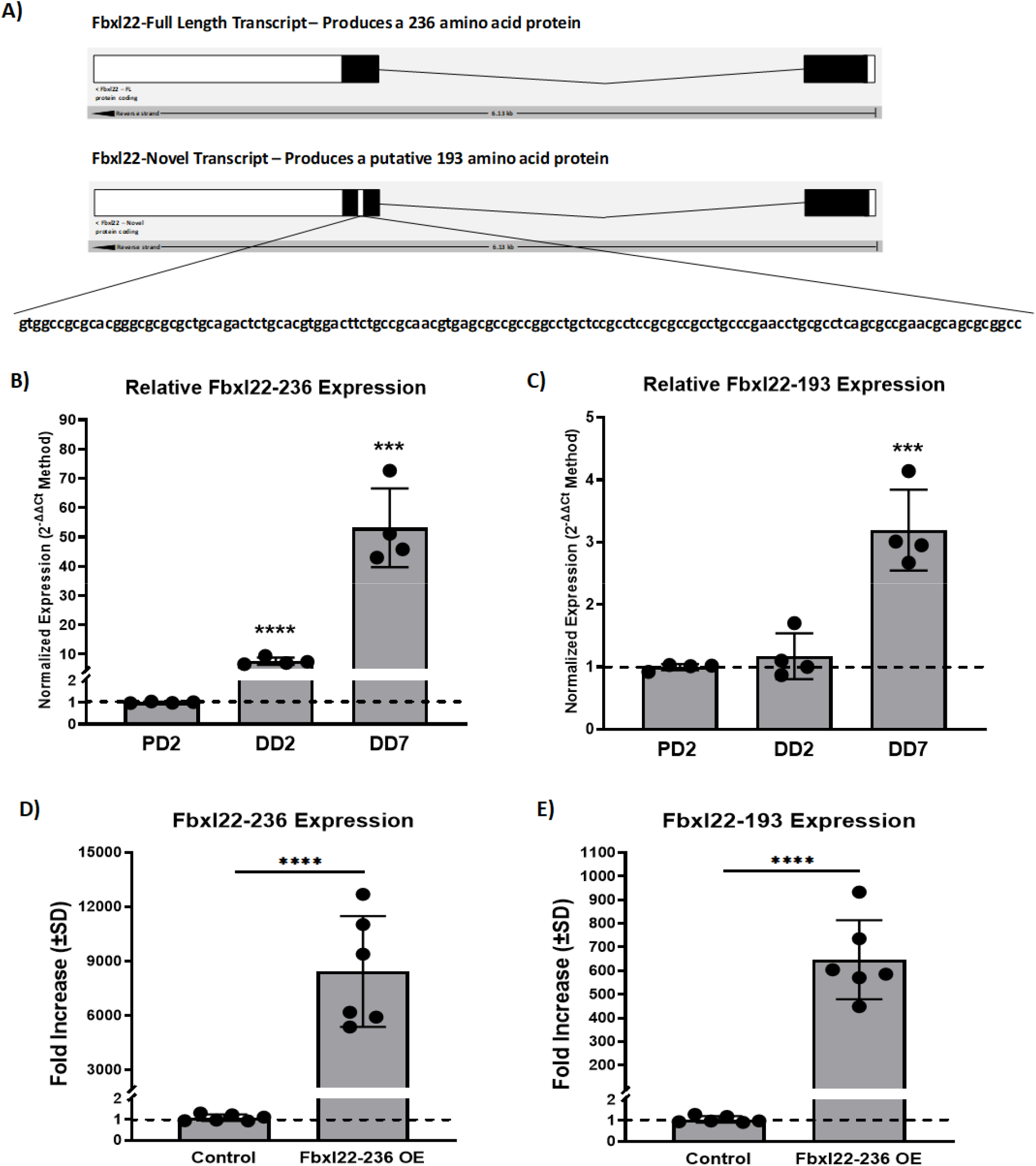
Fbxl22-236 and Fbxl22-193 are expressed in skeletal muscle and are upregulated during muscle cell differentiation. (**A**) Schematic of the full-length (Fbxl22-236) and novel splice variant (Fbxl22-193) transcripts in mouse. Darkened rectangles represent exons containing translated region, open rectangles represent exons containing the untranslated regions and the lines connecting the rectangles represent introns. (**B**) Fbxl22-193 and (**C**) Fbxl22-236 expression increases as C_2_C_12_ cells differentiate. qPCR analysis of Fbxl22 expression in proliferating (PD2) C_2_C_12_ myoblasts and myotubes at early (DD2) and late (DD7) differentiation. RNA was isolated from three biological replicates (n = 3) for each time point and the experiment was repeated at least three times. Significant difference between proliferating and differentiated cells, (*: *P* ≤ 0.01). Ectopic expression of Fbxl22-236 cDNA in C_2_C_12_ cells results in increased levels of both the (**D**) Fbxl22-236 and (**E**) Fbxl2-193 transcripts. Significant difference between controls cells and cells transfected with the Fbxl22-236 expression plasmid, (*: *P* ≤ 0.01).

### Fbxl22 regulatory region is transcriptionally active in muscle cells and is induced by myogenic regulatory factors

The predicted regulatory region of Fbxl22 was cloned using the primer pairs listed in Table 1. The isolated promoter fragments spanned from +40 to −457 and +40 to −967 of the mouse Fbxl22 gene (Fig. 3A). These fragments were fused with the secreted alkaline phosphatase (SEAP) reporter gene to create the pSEAP-Fbxl22-Pro500 and pSEAP-Fbxl22-Pro1000 reporter plasmids. C_2_C_12_ mouse myoblast cells were transiently transfected with the Fbxl22 reporter constructs and SEAP activity levels were analyzed 5-7 days following the switch to differentiation media and compared to SEAP activity levels from cells transfected with the empty pSEAP2-Basic plasmid. The Fbxl22 reporter plasmids showed significant transcriptional activity and had significantly higher expression compared to the empty reporter plasmid alone (Fig. 3A). To determine if the Fbxl22 reporter gene activity mirrors the transcriptional activity of the endogenous gene (Fig. 2B and 2C), the pSEAP-Fbxl22-Pro500 reporter was transfected into C_2_C_12_ myoblasts and media was then analyzed for SEAP activity at 24 hours post-transfection (PD2) and at 2 (DD2) and 7 (DD7) days post-switch to differentiation media. The data shows that the Fbxl22 reporter gene has elevated transcriptional activity as C_2_C_12_ differentiate (Fig. 3B), mirroring the qPCR data.

**Figure 3.**
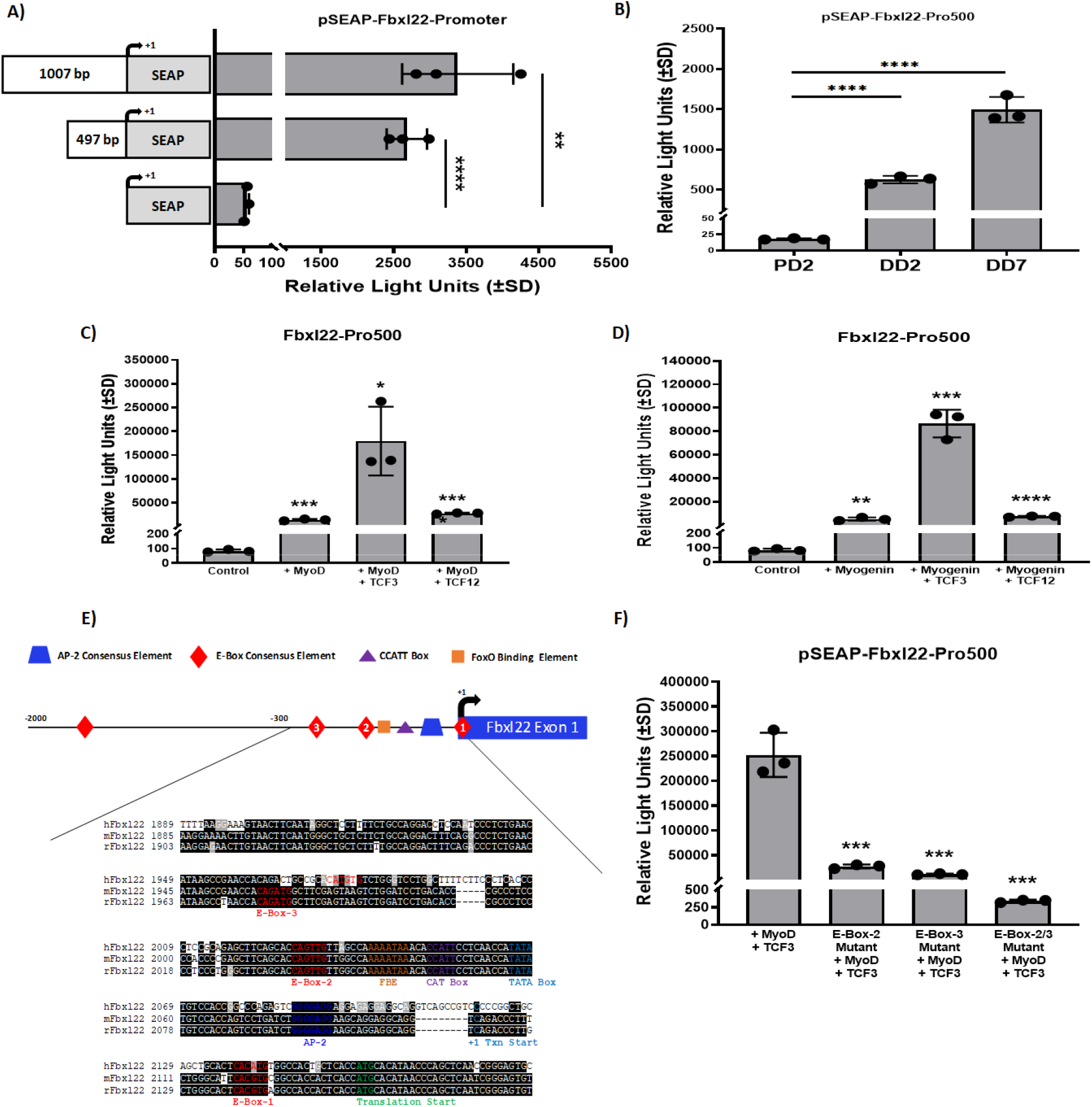
Fbxl22 reporter gene is transcriptionally upregulated during muscle cell differentiation and is activated by myogenic regulatory factors. (**A**) C_2_C_12_ myoblasts were transfected with either an empty reporter plasmid or a reporter plasmid containing ~500 bp or ~1000 bp of the Fbxl22 promoter fused to the Secreted Alkaline Phosphatase (SEAP) reporter gene. Significant differences in activity between the control empty reporter plasmid (pSEAP2-Basic) and the pSEAP-Fbxl22-Pro500 and pSEAP-Fbxl22-Pro1000 reporter constructs (*: *P* ≤ 0.01). C_2_C_12_ myoblasts were transfected with the (**B**) pSEAP-Fbxl22-Pro500 and cells were differentiated for seven days and the media was analyzed at proliferation day 2 (PD2), differentiation day 2 (DD2) and differentiation day 7 (DD7) and SEAP numbers were normalized to β-galactosidase activity to correct for variations in transfection efficiency. Significant differences between the activities of the pSEAP-Fbxl22-Pro500 reporter construct at PD2 compared to DD2 and DD7 (*: *P* < 0.01). C_2_C_12_ myoblasts were transfected with the pSEAP-Fbxl22-Pro500 reporter plasmid alone or in combination with (**C**) MyoD or (**D**) myogenin with or without Tcf3 or Tcf12 expression plasmids. The media was then assayed for SEAP activity at 48 hours following the switch to differentiation media. Significant differences between the activities of the pSEAP-Fbxl22-Pro500 alone and in combination with (**C**) MyoD alone or MyoD±Tcf3 or MyoD±Tcf12 or (**D**) myogenin alone or myogenin±Tcf3 or myogenin±Tcf12 overexpression are indicated (*: *P* < 0.01). (**E**) Analysis of the Fbxl22 promoter region revealed three conserved, putative consensus E-box elements 5’-CANNTG-3’ (red diamonds). N represents any nucleotide. (**F**) C_2_C_12_ myoblasts were transfected with the pSEAP-Fbxl22-Pro500-wild-type, pSEAP-Fbxl22-Pro500-E-box2-mutant, pSEAP-Fbxl22-Pro500-E-box3-mutant, or pSEAP-Fbxl22-Pro500-E-box2/3-mutant reporter constructs with MyoD and Tcf3 expression plasmids. The media was then assayed for SEAP activity at 48 hours following the switch to differentiation media. Significant differences between the activities of the pSEAP-Fbxl22-Pro500-wild-type construct and the pSEAP-Fbxl22-Pro500-E-box-mutant reporter constructs (*: *P* < 0.01). For all Fbxl22 reporter gene experiments, each condition was performed in triplicate and each experiment was repeated at least three times. The graphs are of representative experiments (n = 3) and values correspond to the mean relative light unit (RLU) values ± SD.

Neurogenic atrophy results in a dramatic increase in myogenic regulatory factor (MRF) expression, including significant increases in MyoD1 and myogenin expression (18, 25, 44, 48, 63). Therefore, since MyoD1 and myogenin are elevated during neurogenic atrophy and are capable of binding canonical E-box elements in muscle-specific genes, we evaluated the ability of the Fbxl22 reporters to respond to ectopic expression of MyoD1 and myogenin in C_2_C_12_ cells transiently transfected with the Fbxl22-Pro500 wild-type reporter constructs. Cells transiently transfected with MyoD1 or myogenin in combination with Tcf3 or Tcf12 expression plasmids showed significantly higher Fbxl22 reporter gene activity compared to cells that were not (Fig. 3C and 3D).

### Identification of conserved, putative E-box elements in the Fbxl22 regulatory region

To better assess the level of conservation between the regulatory regions of mouse, rat, and human Fbxl22 and to identify potential transcriptional regulatory elements, the genomic sequences corresponding to the regulatory region of this gene for mouse, rat, and human were aligned (Fig. 3E). Several putative enhancer elements were identified in the proximal promoter region of Fbxl22, including one putative AP-2 consensus sequence (37), one putative CCAAT box (43), and one putative FoxO binding element (56, 66) (Fig. 3E). Furthermore, since MRFs are elevated during neurogenic atrophy, are capable of binding E-box elements in muscle-specific genes, and appear to positively regulate Fbxl22 reporter gene activity, we analyzed the Fbxl22 regulatory region and identified two putative E-box elements located just upstream of the start of transcription of the Fbxl22 gene. These two putative E-box elements lie between −145 and −70 of the proximal promoter region of Fbxl22 (Fig. 3E). To determine if the E-box elements are necessary for proper Fbxl22 promoter activity, site-directed mutagenesis was performed using the −500 bp Fbxl22 reporter plasmid, resulting in the generation of the pSEAP-Fbxl22-Pro500 E-box2, pSEAP-Fbxl22-Pro500 E-box3, and the pSEAP-Fbxl22-Pro500 E-box2/3 mutant constructs. Mutation of either E-box in the 500 bp Fbxl22 promoter construct resulted in significantly lower reporter gene activity compared to the wild-type reporter construct in transiently transfected C_2_C_12_ cells (Fig. 3F), suggesting that MRF activation of Fbxl22 expression may occur via these proximal E-box elements.

### Overexpression of the Fbxl22 isoforms in unchallenged TA muscles for 7, 14 and 28 days post electroporation

We sought to manipulate gene expression levels of both Fbxl22 isoforms (−236 and −193) though an in vivo electroporation approach into unchallenged TA muscles for periods of 7,14, and 28 days. Transfection of the full-length Fbxl22-236 construct in the TA resulted in no changes in muscle mass at 7 and 28 days, but an increase after 14 days post electroporation (P = 0.01; Fig. 4C). Measurement of muscle fiber CSA showed no differences between Fbxl22-236 transfected muscles and EV control muscles after 7 days. A shift in the distribution of fiber CSAs towards smaller muscle fibers was observed in Fbxl22-236 transfected muscles after 14 days but was no longer evident after 28 days (Fig. 4D). On the other hand, transfection of the novel Fbxl22-193 isoform in the TA lead to a significant increase in muscle mass after 7 days, but no differences in muscle mass at 14 or 28 days (Fig. 4E). In terms of muscle CSA, there were increases in the percentage of smaller and larger fibers with the presence of Fbxl22-193 overexpression compared to EV control muscles at 14 and 28 days (Fig. 4F).

**Figure 4.**
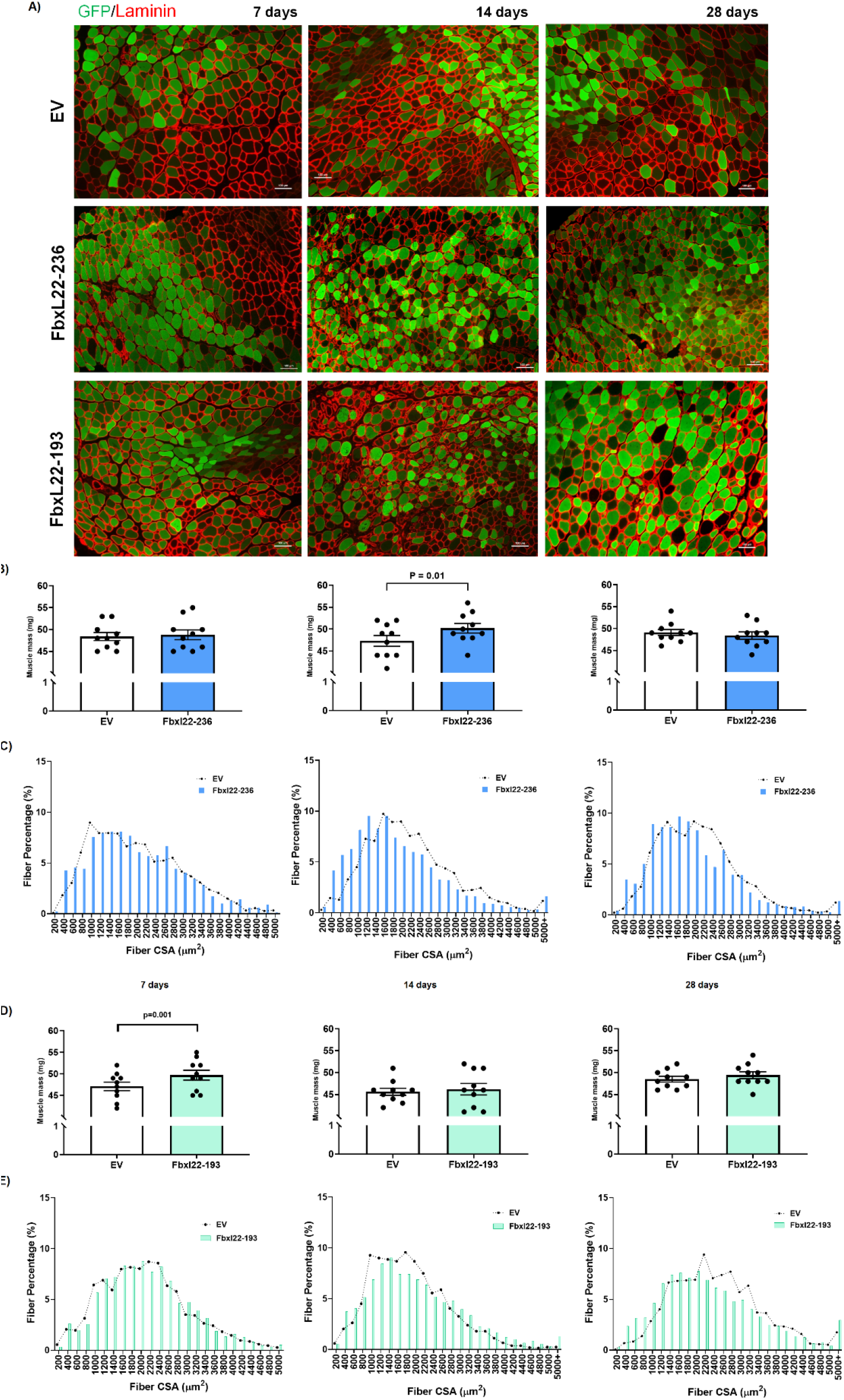
Overexpression of Fbxl22 isoforms in unchallenged tibialis anterior (TA) muscle for 7, 14 and 28 days. (**A**) Representative microscope images for GFP fluorescence and laminin in TA mouse muscles transfected with empty vector (EV), Fbxl22-236 and −193 plasmids (10x magnification; scale bar = 100μm) highlighting transfection efficiency and muscle morphology after 7 (left), 14 (middle) and 28 (right) days. Muscle mass measurements (milligrams) for muscles transfected with either the EV or Fbxl22-236 (**C**) and −193 (**E**) plasmids (n=10/group) after 7 (left), 14 (middle) and 28 (right) days. Muscle cross-sectional area (CSA) distribution for EV and Fbxl22-236 (**D**) and −193 (**F**) transfected muscles (7 days = left; 14 days = middle; 28 days = right). GFP-positive fibers were measured for CSA, with ≥ 350 transfected fibers analyzed per animal, per muscle (n= 6/group) and data presented as a percentage of sizes between 0-5000μm. For 14- and 28-day muscle CSA data, fibers that were 5000μm-plus were included in the analysis due to the phenotype observed with Fbxl22-236 and −193 overexpression at these time points. Data presented as Mean ± SEM.

At 14 days, Fbxl22-236 and Fbxl22-193 transfected muscles displayed increased cell infiltration, necrosis and degeneration, and the large fibers were rounded in appearance which is abnormal (Fig. 5A). With prolonged Fbxl22-193 overexpression for 28 days, we observed large fibers with numerous centrally located nuclei and reduced amounts of cell infiltration (Fig. 5A) suggestive of regeneration. Overexpression of −236 and −193 isoforms resulted in a significant amount of transfected fibers with centrally located nuclei after 14 and 28 days (Fig 5B and 5C).

**Figure 5.**
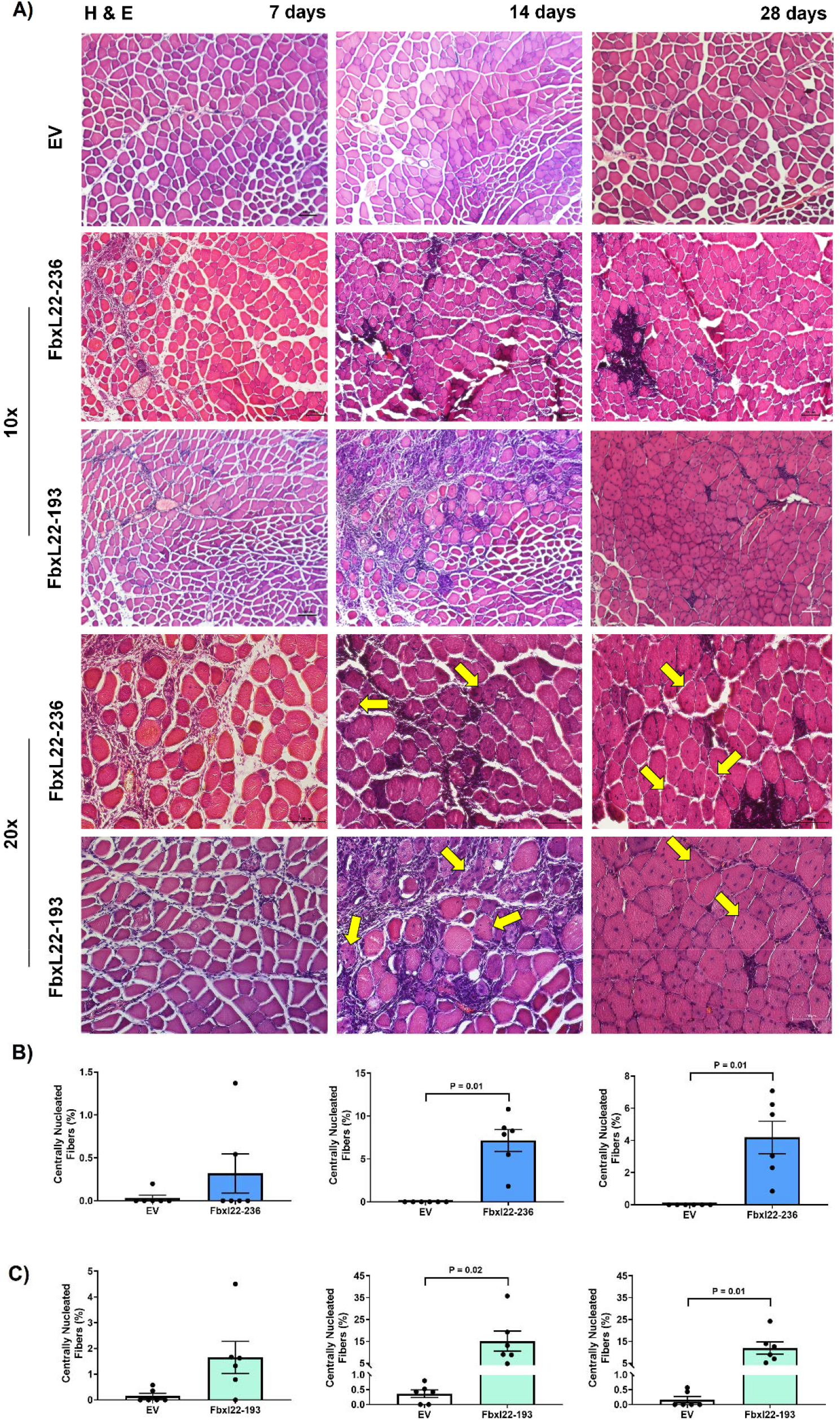
Quantification of Centrally located nuclei in tibialis anterior (TA) muscle fibers with Fbxl22 isoform overexpression for 7, 14 and 28 days. Representative microscope images for hematoxylin and eosin (H & E) in TA mouse muscles transfected with empty vector (EV), Fbxl22-236 and −193 plasmids (10x and 20x magnification; scale bar = 100μm) highlighting muscle morphology after 7 (left), 14 (middle) and 28 (right) days. Examples of centrally located nuclei are identified on H & E images with yellow arrows. Fibers with centrally located nuclei with quantified for −236 (B) and −193 (C) transfected TA muscles for 7 (left), 14 (middle) and 28 (right) days (n = 5/group). Percentage of centrally nucleated myofibers was calculated by dividing the number of transfected fibers containing ≥1 centrally located nuclei by the total number of transfected fibers. Statistical significance is depicted where present. Data presented as Mean ± SEM.

At the biochemical level, there was no change in total ubiquitin or p62 levels with Fbxl22-236 overexpression at 14 days, but there was an increase in LC3B II content (Fig. 6B). Further, a reduction in dystrophin protein and significant elevations in desmin and vimentin were observed (Fig. 6C). Overexpression of Fbxl22-236 for 7 days resulted in alterations in total ubiquitin, p62 and LC3B II protein content, but no significant changes in dystrophin and intermediate filament protein levels (Fig. S2). The only significant differences observed in sarcomeric protein levels after 28 days of Fbxl22-236 overexpression was a decrease in α-actinin and an increase in vimentin (Fig. S2). In comparison, there were significant increases in total ubiquitin, p62 and LC3B II content in Fbxl22-193 transfected muscles compared to the EV control group at 14 days (Fig. 6E). In addition, we observed a significant reduction in dystrophin protein and changes in α-actinin isoform levels (Fig. 6F). The alterations in α-actinin isoforms were accompanied by 3-to 4-fold increases in desmin and vimentin (Fig. 6F). After 7 days of electroporation, Fbxl22-193 transfected muscles had elevation in p62 levels, reduction in dystrophin, and alterations in α-actinin isoforms and vimentin (Fig. S3). At 28 days, total ubiquitin and LC3B II content remained elevated (1.5-fold), while no significant changes in cytoskeleton protein content were observed (Fig. S3). The overall changes in key sarcomeric proteins may be indicative of destabilization in the myofilament lattice and the appearance of regenerating muscle fibers (13, 26, 34, 41).

**Figure 6.**
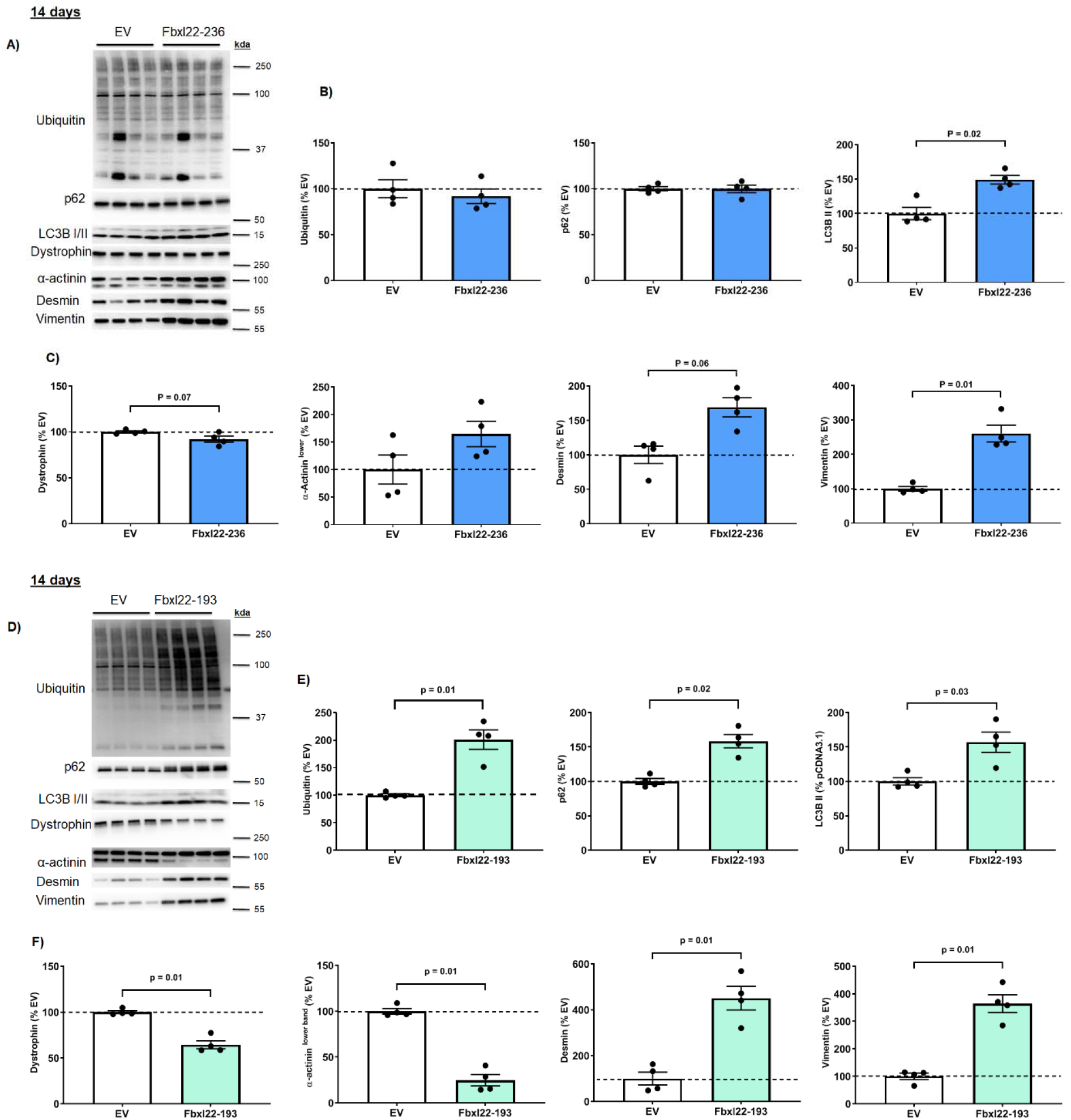
Alterations in ubiquitin, autophagy markers and cytoskeleton protein levels in Fbxl22 transfected tibialis anterior (TA) muscles after 14 days. (**A and D**) Representative immunoblot images for empty vector (EV), Fbxl22-236 and −193 transfected muscles after 14 days (n=4/group). (**B and E**) Total protein levels for ubiquitin, p62 and LC3B II. (**C and F**) Total protein levels for dystrophin, α-actinin (lower band), desmin, and vimentin. Total protein loading was used as the normalization control for all blots. Statistical significance is depicted where present. Data presented as Mean ± SEM.

### Overexpression of Fbxl22 isoforms in unchallenged lateral and medial gastrocnemius muscles for 14 days

The H&E staining of both GA muscles showed a similar pattern to the TA muscle after 14 days of overexpression, where GA muscles transfected with either Fbxl22 isoform displayed increased cell infiltration, necrosis and degeneration of skeletal muscle fibers (Fig. 7A and 7B). In the lateral GA muscle, the overexpression of Fbxl22-236 led to a significant increase in muscle mass (Fig. 7C), while overexpression of the Fbxl22-193 isoform caused no change in muscle mass compared to EV transfected muscles (Fig. 7D). In Fbxl22-236 transfected muscles, there was a shift in the distribution of fiber CSA primarily towards smaller fibers compared to the EV control muscle (Fig. 7C). Enlarged fibers could be found in the Fbxl22-236 transfected muscles, however, they were usually abnormally rounded in shape. The Fbxl22-193 transfected muscle displayed a slight shift towards an increase in smaller fibers compared to the contralateral EV control group (Fig. 7D). In the medial GA muscle, Fbxl22-236 transfected muscles displayed a significant reduction in muscle mass and a leftward shift towards smaller muscle fibers compared to the EV transfected muscles (Fig. 7E). The overexpression of Fbxl22-193 isoform in the medial GA muscle caused no change in muscle mass but did produce a shift in the distribution of fiber CSA primarily towards an increase in the number of smaller muscle fibers (Fig. 7F). In addition, we observed a significant increase in fibers with centrally located nuclei for −193 transfected GA muscles after 14 days (Fig 7D and 7F).

**Figure 7.**
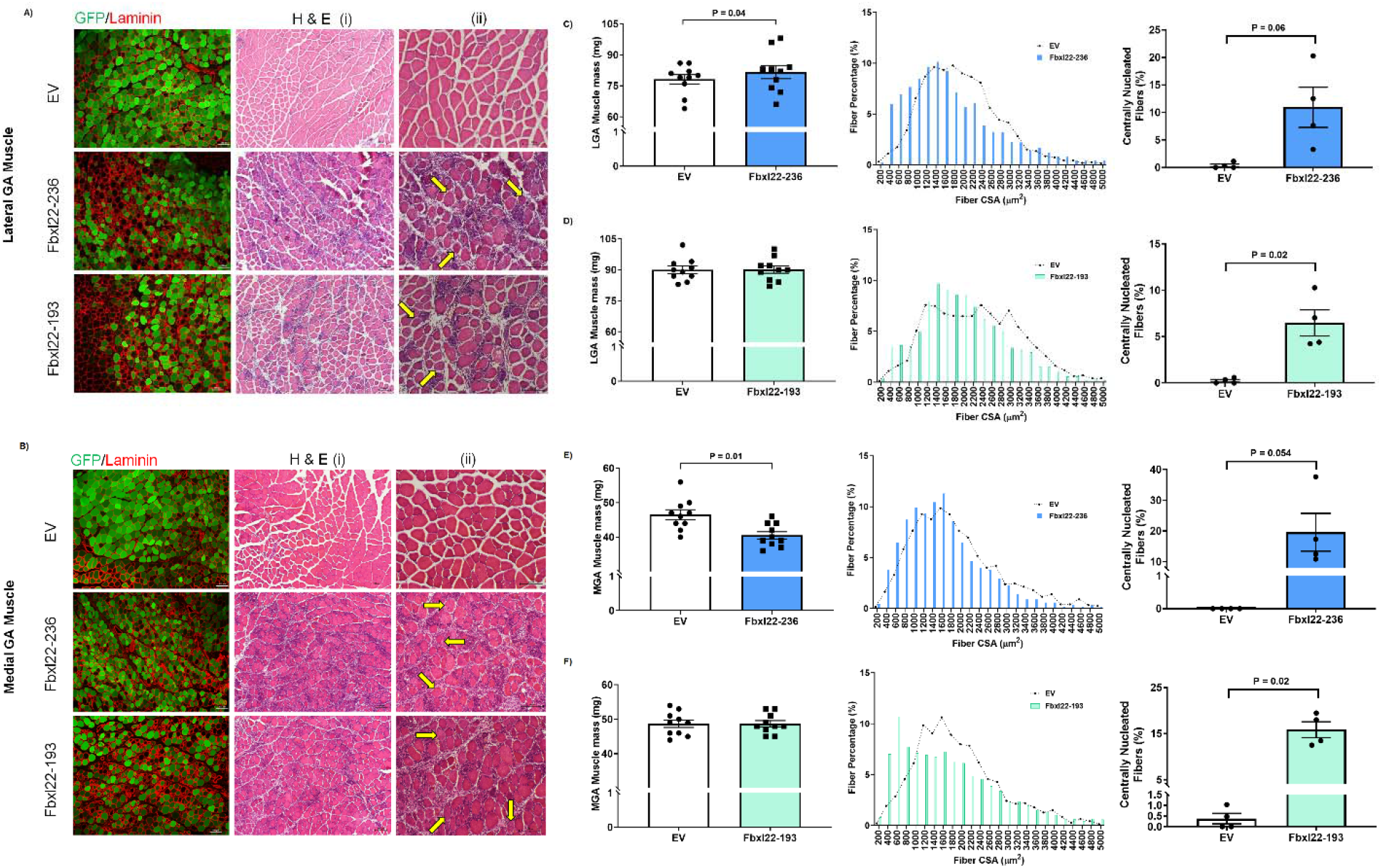
Overexpression of Fbxl22 isoforms in unchallenged lateral and medial gastrocnemius muscles for 14 days. Representative microscope images for GFP fluorescence, laminin and H&E in lateral (LGA; Panel **A**) and medial (MGA; Panel **B**) gastrocnemius mouse muscles transfected with empty vector (EV) or Fblx22-236 and Fbxl22-193 constructs (i, 10x magnification and ii, 20x magnification; scale bar = 100μm) highlighting transfection efficiency and muscle morphology. Examples of centrally located nuclei are depicted on H & E images with yellow arrows. Muscle mass measurements (milligrams) for LGA (Panel **C** and **D**) and MGA (Panel **E** and **F**) muscles transfected with either the EV, Fbxl22-236 and Fbxl22-193 constructs (n=10/group). Muscle cross-sectional area (CSA) distribution in LGA (Panel **C** and **D**) and MGA (Panel **E** and **F**) transfected muscles with either the EV, Fbxl22-236 or Fbxl22-193 constructs. GFP-positive fibers were measured for CSA, with ≥ 350 transfected fibers analyzed per animal, per muscle (n= 5/group) and data presented as a percentage of sizes between 0-5000μm. Fibers with centrally located nuclei (≥1) quantified in LGA (Panel **C** and **D**) and MGA (Panel **E** and **F**) transfected muscles with either the EV, Fbxl22-236 and Fbxl22-193 constructs (n=4/group). Statistical significance is depicted where present. Data presented as Mean ± SEM.

The histological changes observed with Fbxl22 overexpression in the GA muscle corresponded with increased levels of markers of protein degradation (total ubiquitin, p62 and LC3B II) and alterations in cytoskeleton proteins (dystrophin, α-actinin, desmin, and vimentin) compared to EV transfected muscle (Fig. S4 and S5). These observations were present in both the lateral and medial GA muscles after overexpression of the full-length and novel Fbxl22 isoforms for 14 days. These data highlight the variation in the morphological and biochemical response of different skeletal muscles (TA versus gastrocnemius), depending on which Fbxl22 isoform is overexpressed.

### Knockdown of Fbxl22 leads to partial muscle sparing in wild type muscle during denervation-induced muscle atrophy

Given the early upregulation of Fbxl22 expression following denervation, we studied whether knock down of Fbxl22 in muscle would have an effect on muscle atrophy following denervation. We first confirmed that our Fbxl22 RNAi construct could knock down Fbxl22 mRNA expression after 3 days of denervation. The Fbxl22 RNAi transfected muscles displayed a 60-70% reduction in endogenous expression of Fbxl22 mRNA (for both isoforms) in the LGA and MGA muscles compared to the EV transfected muscles (Fig. S6). After 7 days of denervation, total GA mass in WT mice significantly decreased in EV transfected and Fbxl22 RNAi transfected muscles, with no significant difference between groups (Fig. 8A). In relation to the contralateral control muscle, EV transfected muscles exhibited a 31% loss in muscle mass, whereas Fbxl22 RNAi transfected muscles displayed a 27% loss (P = 0.054; Fig. 8B). When we examined the two heads of GA separately, we found that in the LGA muscle there was no muscle sparing in the Fbxl22 RNAi transfected group (Fig. 8C), but in the MGA there was significant muscle sparing in the presence of the Fbxl22 RNAi (P = 0.03; Fig. 8D). The mean fiber CSA for GFP-positive fibers was significantly larger in the RNAi transfected muscles compared to the EV group (P = 0.04; Fig. 8E). Measurement of fiber CSA in both heads of the GA revealed that Fbxl22 RNAi resulted in partial preservation of muscle CSA size compared to non-transfected fibers, as signified by a reduced leftward shift in distribution towards smaller muscle fibers (Fig. 8F). In comparison, the EV displayed a leftward shift towards smaller muscle fibers that was similar to the non-transfected fibers (Fig. 8G).

**Figure 8.**
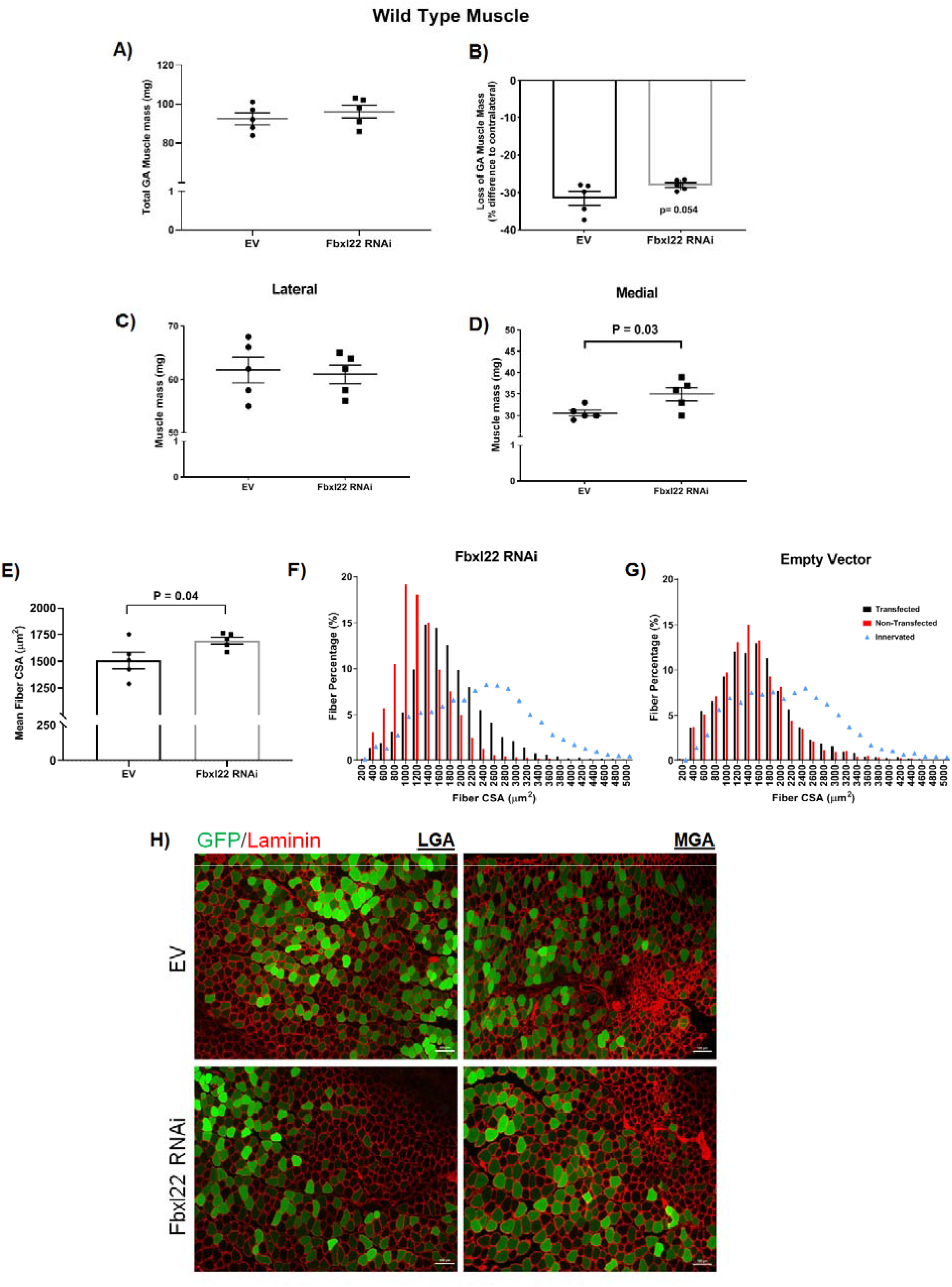
Fbxl22 knockdown leads to partial muscle sparing after 7 days of denervation in wild type mice. (**A**) Total muscle mass for gastrocnemius (GA; lateral and medial) muscles transfected with either empty vector (EV) or Fblx22 RNAi constructs. (**B**) Loss of GA muscle mass with either EV or Fbxl22 RNAi present compared to contralateral control (innervated) muscle. Data expressed as percent lost compared to the contralateral control. Muscle mass for **(C)** lateral and **(D)** medial gastrocnemius muscles transfected with EV or Fbxl22 RNAi construct after 7 days of denervation. (**E**) Mean Fiber cross-sectional area (CSA) for GFP-positive only fibers in transfected GA muscles was determined. Muscle cross-sectional area (CSA) distribution in transfected muscles with the **(F)** Fbxl22 RNAi or **(G)** EV constructs. GFP-positive fibers were measured for CSA, with ≥ 350 transfected fibers analyzed per animal, per muscle (n=5/group). In addition, the muscle CSA distributions for non-transfected (red bars) and innervated (blue triangle) contralateral control muscles were measured. Data presented as a percentage of sizes between 0-5000μm. (**H**) Representative images for GFP fluorescence and laminin stain in Fbxl22 RNAi and EV transfected lateral and medial gastrocnemius muscles after 7 days of denervation (10x magnification; scale bar = 100μm). Statistical significance is depicted where present. Data presented as Mean ± SEM (N= 5/group).

### Knockdown of Fbxl22 in MuRF1 knockout muscle results in greater muscle sparing under denervation-induced muscle atrophy

As muscle sparing is evident in the MuRF1 KO mice at 14 days, but not 7 days (9), we sought to determine the effect of knockdown of Fbxl22 expression in MuRF1 KO muscles. After 7 days of denervation in MuRF1 KO mice, total GA mass was significantly smaller in the EV transfected muscles compared to the Fbxl22 RNAi transfected muscle (P = 0.03; Fig. 9A). MuRF1 KO muscles transfected with the EV exhibited a 27% loss in muscle mass compared to the contralateral control (Fig. 9B), while MuRF1 KO muscles transfected with the Fbxl22 RNAi construct showed significant muscle sparing with only a 19% loss in muscle mass compared to the contralateral control muscle (P = 0.01; Fig. 9B). Examination of the two heads separately showed that in the LGA muscle there was partial (albeit non-significant) sparing in the Fbxl22 RNAi transfected group (Fig. 9C) and significant muscle sparing in the MGA muscle with the presence of the Fbxl22 RNAi (P = 0.01; Fig. 9D). The mean fiber CSA for GFP-positive fibers was significantly larger in the RNAi transfected muscles compared to the EV group (P = 0.01; Fig. 9E). Analysis of muscle fiber CSA revealed Fbxl22 RNAi resulted in preservation of muscle CSA size compared to non-transfected fibers, as signified by a lack of a leftward shift in distribution towards smaller muscle fibers (Fig. 9F). In comparison, the EV transfected fibers displayed a leftward shift towards smaller CSAs that was similar to the non-transfected fibers (Fig. 9G).

**Figure 9.**
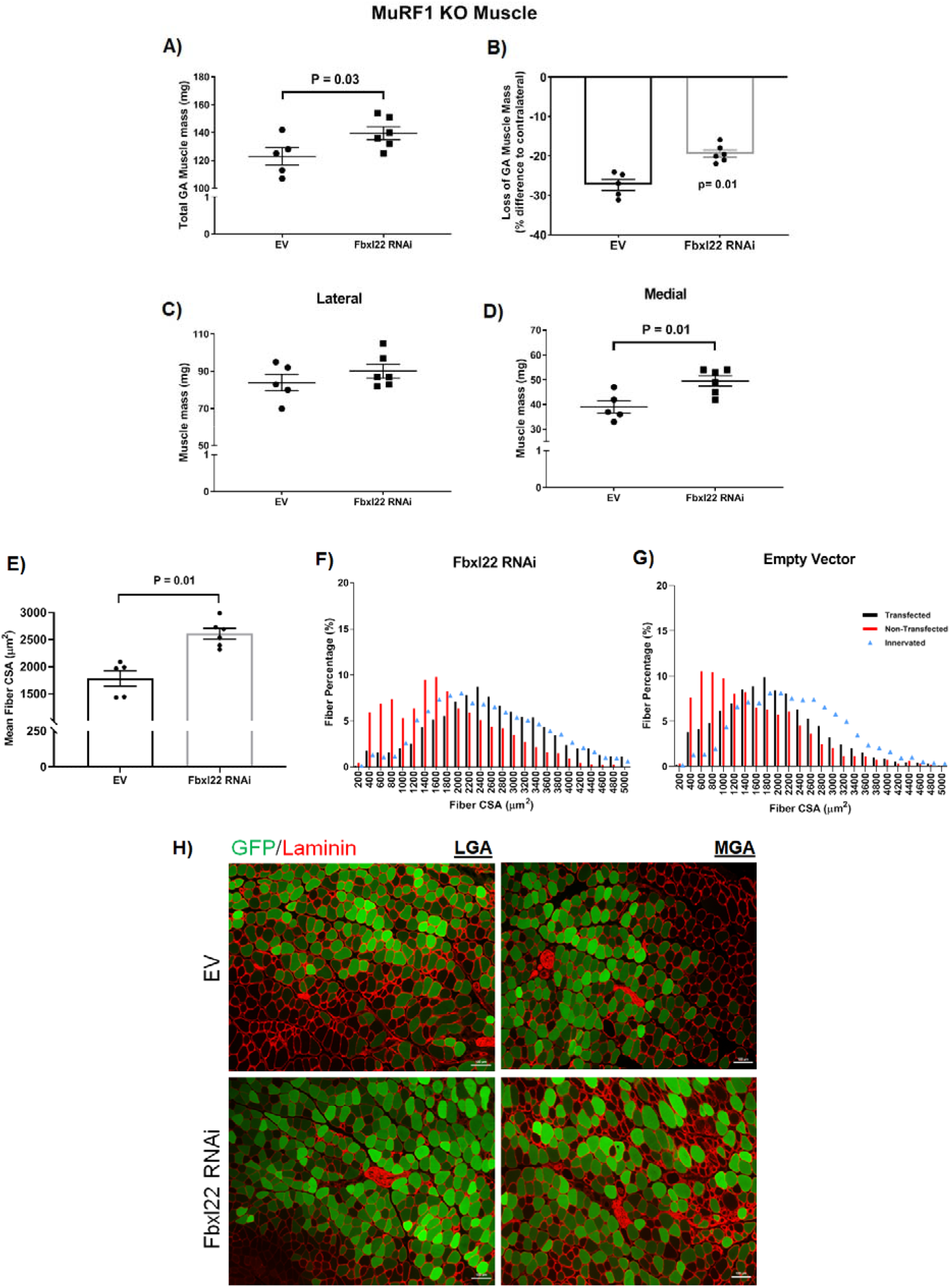
Knockdown of Fbxl22 in MuRF1 KO muscle results in improved muscle sparing after 7 days of denervation. (**A**) Total muscle mass for gastrocnemius (GA; lateral and medial) muscles transfected with either empty vector (EV) or Fblx22 RNAi constructs. (**B**) Loss of GA muscle mass with either EV or Fbxl22 RNAi present compared to contralateral control (innervated) muscle. Data expressed as percent loss compared to contralateral control. Muscle mass for **(C)** lateral and **(D)** medial gastrocnemius muscles transfected with EV or Fbxl22 RNAi construct after 7 days of denervation. (**E**) Mean Fiber cross-sectional area (CSA) for GFP-positive only fibers in transfected GA muscles was determined. Muscle CSA distribution in transfected muscles with the **(F)** Fbxl22 RNAi or **(G)** EV constructs. GFP-positive fibers were measured for CSA, with ≥ 350 transfected fibers analyzed per animal, per muscle (n= 5/group). In addition, the muscle CSA distributions for non-transfected (red bars) and innervated (blue triangle) contralateral control muscles were measured. Data presented as a percentage of sizes between 0-5000μm. (**H**) Representative images for GFP fluorescence and laminin stain in Fbxl22 RNAi and EV transfected lateral and medial gastrocnemius muscles after 7 days of denervation (10x magnification; scale bar = 100μm). Statistical significance is depicted where present. Data presented as Mean ± SEM (N= 5-6/group).

## Discussion

Skeletal muscle has the ability to regulate its size in response to internal and external cues (such as neural activity, nutritional status, etc.) throughout the lifespan (60). Homeostasis of skeletal muscle size occurs through a balance between protein synthesis and breakdown, with the balance shifting to net protein breakdown during skeletal muscle atrophy events (7, 47, 58). The UPS is an integral part of the protein breakdown process in skeletal muscle and E3 ubiquitin ligases are critical in targeting substrates for ubiquitination (8, 55). Surprisingly, to date only a handful of E3 ubiquitin ligases have been identified and characterized as important in the process of skeletal muscle atrophy (9, 28, 45, 51, 57). In our present study, we have identified Fbxl22 and a novel splice variant that code for E3 ubiquitin ligases that are significantly upregulated in skeletal muscle exposed to neural inactivity. From our previously published microarray data (25), we found the RIKEN gene 1110004B15RIK to be at the top of the list of genes upregulated after 3 days of denervation in wild type and MuRF1 KO muscle. We subsequently identified this RIKEN gene to be Fbxl22. Through an in-depth time course analysis of denervation, Fbxl22 isoforms were found to significantly increase early in the atrophy process and return to baseline levels by 7 days. In comparison, MuRF1 and MAFbx expression remain elevated for a longer time period in neurogenic muscle atrophy models, returning to baseline after 14 days (9, 25, 38, 48, 59). Thus, we hypothesized that Fbxl22 could be involved in the early initiation of the muscle atrophy process.

Fbxl22 is a member of the F-box family (which includes MAFbx, MUSA1, and SMART (9, 45, 57)) and has previously been observed to play a role in sarcomeric turnover in cardiac muscle (62). To our knowledge, the study by Spaich and colleagues (62) is the only other published investigation into the role of Fbxl22 in striated muscle. In the study by Spaich et al, the authors observed localization of Fbxl22 at the sarcomeric z-line and *in vitro* overexpression studies identified filamin C and α-actinin to be Fbxl22 targeted substrates for ubiquitination. Through analysis of Fbxl22 overexpression at multiple time points and in multiple muscles, we observed significant alterations in α-actinin isoforms, desmin, and vimentin which are all integral for maintaining muscle fiber integrity and function (26, 34, 40, 50). Specifically, both Fbxl22 isoforms cause a reduction in one of the α-actinin isoforms and significant elevations in desmin and vimentin. Interestingly, expression of desmin and vimentin is suggested to be indicative of regenerating muscle fibers (13, 26, 65). It is important to note that desmin has been observed to be targeted for degradation in muscle atrophy models, specifically by the E3 ubiquitin ligase TRIM32 (17). We also observed significant reductions in dystrophin protein content in Fbxl22 transfected muscles. Dystrophin is essential for lateral force transmission and protecting skeletal muscle from contraction-induced injury (32, 33, 41, 54). The alterations in these sarcomeric proteins coincided with the development of a degenerative muscle fiber phenotype in Fbxl22 transfected tissues. Other E3 ubiquitin ligases, such as MuRF1, have been reported to target a distinct subset of sarcomeric proteins (such as titin and myosin heavy chain) for degradation in skeletal muscle (14, 15, 17, 53), suggesting that the protein turnover that accompanies muscle atrophy is likely orchestrated by several different E3 ligases.

Upon neurogenic atrophy, there is a dramatic increase in MRF expression including MyoD1 and myogenin (18, 44, 48). Within the Fbxl22 promoter, we identified two putative E-box elements immediately upstream of the start of transcription, of which one is fully conserved between rodents and humans. Furthermore, Fbxl22 reporter gene activity appears to be positively regulated by MRF expression, and thus highlights the potential role for Fbxl22 in the early initiation of the muscle atrophy process. Interestingly, the development of a degenerative fiber phenotype in our overexpression studies took a progressive period of time to develop (up to 14 days). The prolonged expression of Fbxl22 alone may explain the appearance of the observed myopathy, since Fbxl22 induction appears to be short-lived and returns rapidly to baseline after just 3 days of denervation. Furthermore, it is also possible that other genes that are differentially expressed in response to denervation may work to direct the activity of Fbxl22 during skeletal muscle atrophy and prevent the development of myopathy (25). Although, we observed no dramatic effect on muscle mass with Fbxl22 isoform overexpression, except in the MGA muscle, we did observe a distinct phenotype in muscle fiber CSA with the appearance of small degenerating fibers and large abnormal fibers which may have masked the lack of effect on overall muscle mass. In the MGA muscle, the fiber CSA distribution shifted towards the left due to an increase in smaller fibers with no large abnormal fibers present, which coincided with a significant loss of muscle mass with Fbxl22 overexpression. The lack of uniform response on muscle mass in Fbxl22 isoform overexpression studies highlights the complexity of studying a single component of the muscle atrophy program in multiple hindlimb muscles (3, 6, 21).

The manifestation of the degenerative phenotype in Fbxl22 transfected muscles appears to differ between muscles and is dependent on which Fbxl22 isoform is overexpressed, this being reflected in the subtle differences observed in our biochemical analyses. We have previously observed muscle-specific changes in levels of atrophy upon implementation of a hindlimb unloading model (2, 3, 46). While differences in muscle fiber type, neural input, and/or loading history may all contribute to our previous findings, it is important to highlight that the changes in E3 ubiquitin ligase expression (MuRF1 and MAFbx) varied between the rat soleus, medial gastrocnemius and TA muscles during the unloading period (3). It would appear that different muscles may atrophy in different ways at the molecular level depending on which genes are and are not expressed (1, 3, 25). To note, we only implemented a neural inactivity stimulus for inducing atrophy and thus future studies are needed to explore Fbxl22 expression in other atrophy models where the motor unit remains intact (e.g. hindlimb suspension, glucocorticoids). It is possible that the action of Fbxl22 may differ between hindlimb muscles and this may explain why we observed muscle atrophy in MGA transfected muscles, but a degenerative myopathy in LGA and TA transfected muscles. In addition, differences in the ratio of E1 and E2 enzymes present in each manipulated muscle (TA, LGA and MGA) and/or differences in Fbxl22 E2 binding partners could influence the muscles’ response to Fbxl22 overexpression. Furthermore, we observed different levels of total ubiquitin between Fbxl22 isoforms and the muscle transfected, which may indicate different substrates being targeted for degradation. Future studies will seek to identify which E2 enzymes pair with Fbxl22 and the specific substrates that Fbxl22 targets for ubiquitination.

In the reported literature, Fbxl22 has been indirectly assessed in skeletal muscle through transcriptomic analysis and genome-wide screens in various models. In human skeletal muscle exposed to either bed rest or unilateral lower limb immobilization, Fbxl22 does not appear to be transcriptionally active in vastus lateralis biopsy samples between 2 and 14 days of intervention (52). Interestingly, Fbxl22 has been reported in C_2_C_12_ muscle cells to inhibit SMAD4 translocation and appears to modulate transforming-growth factor-β1 (TGFβ1) signaling (20). To note, our previously published microarray data and other studies have independently observed alterations in the TGF-β signaling pathway in denervated muscle tissues (10, 25). As the network of TGF-β ligands have been observed to be negative regulators of muscle mass (11, 12), future studies may seek to explore the possible molecular interactions between Fbxl22 and TGF-β signaling in muscle mass regulation.

Multiple E3 ubiquitin ligases have been targeted individually for knock down or knock out in various skeletal muscle atrophy models, often resulting in sparing of muscle mass (9, 28, 45, 51, 57). We report that RNAi knockdown of Fbxl22 mRNA expression leads to partial muscle sparing in wild type muscle undergoing denervation-induced atrophy. However, knock down of Fbxl22 in muscle with a deletion of MuRF1 resulted in significantly greater sparing of muscle mass following 7 days of denervation. This result is particularly interesting since the deletion of MuRF1 alone does not lead to sparing of muscle mass at 7 days of denervation. Interestingly, MuRF1 KO muscles display increased Fbxl22 expression after 3 days of denervation, and thus Fbxl22 could be an important factor driving the onset of the early atrophy phase, prior to when the deletion of MuRF1 results in muscle sparing (9, 25). In both wild type and MuRF1 KO muscle, the medial gastrocnemius muscle displayed significant muscle sparing with the presence of the Fbxl22 RNAi vector compared to the lateral muscle after 7 days of denervation. In addition, the medial gastrocnemius muscle was the only muscle to display muscle atrophy with Fbxl22 overexpression. MuscleDB database analysis shows varying basal mRNA expression levels of Fbxl22 (64) in various hindlimb muscles (plantaris, TA, gastrocnemius, etc.), which may explain the differing degrees of sparing observed during neurogenic muscle atrophy. Another possible explanation is the difference in transfection efficiency for the plasmid vectors between the lateral and medial gastrocnemius, which differ in size and fiber type (61). Indeed, we observed significant effects of muscle sparing with the Fbxl22 RNAi when single transfected muscle fibers were analyzed for CSA verses whole muscle mass measures which includes non-transfected fibers. The development of a muscle specific Fbxl22 knockout animal or use of an adeno-associated viral vector (AAV) may be advantageous for future studies (19, 49). Finally, subsequent studies are required to explore the possible mechanisms for the strong effect in muscle sparing with combined inhibition of Fbxl22 and MuRF1 upon neurogenic muscle atrophy.

## Conclusions

In the present study we extend our understanding of Fbxl22 by identifying a novel splice variant of the gene, by demonstrating that in vivo overexpression causes muscle atrophy and possibly a myopathy after 14 days, and that knockdown of Fbxl22 leads to muscle sparing in a neurogenic muscle atrophy model. Fbxl22 overexpression resulted in increased protein degradation and alterations in key sarcomeric proteins that maintain muscle fiber structure and function. Also, by targeting two E3 ubiquitin ligases (Fbxl22 and MuRF1) simultaneously, we observed enhanced muscle sparing after only 7 days of exposure to an atrophy stimulus. Future studies will seek to investigate the potential role of Fbxl22 in various other atrophy stimuli such as disuse, nutritional deprivation and glucocorticoid treatment.

## Abbreviations

AAV: adeno-associated viral vector;
CSPD: 3-(1-chloro-3′-methoxyspiro-(adamantane-4,4′-dioxetane)-3′-yl) phenyl dihydrogen phosphate;
DMEM: Dulbecco’s Modified Eagle’s Medium;
Fbxl22: F-box and leucine-rich protein 22;
GEO: Gene Expression Omnibus;
GFP: Green Fluorescent Protein;
H&E: Hematoxylin & Eosin;
LGA: lateral gastrocnemius;
MAFbx: muscle atrophy F-box;
MGA: medial gastrocnemius;
MuRF1: muscle-specific RING finger 1;
MUSA1: muscle ubiquitin ligase of the SCF complex in atrophy-1;
MRF: myogenic regulatory factor;
RLU: Relative Light Units;
SEAP: Secreted Alkaline Phosphatase;
SMART: Specific of Muscle Atrophy and Regulated by Transcription;
TA: tibialis anterior;
TGFβ1: transforming-growth factor-β1;
UBR5: ubiquitin protein ligase E3 component n-recognin 5;
UPS: ubiquitin-proteasome system.

## Data Availability

The data that support the findings of this study are available from the corresponding author upon reasonable request.

## Online supplementary material

Additional supporting information may be found online via the link for Figshare data: https://figshare.com/s/4bae1a93d95da33645d3.

## Author Contributions

Conception and design of the experiments: DCH, LMB, DSW and SCB. Collection, analysis and interpretation of data: DCH, LMB, JRD, SAL, DSW and SCB. Drafting the article or revising it critically for important intellectual content: DCH, LMB, DSW and SCB. All authors read and approved the final version of the manuscript and all authors listed qualify for authorship.

