## Supplemental Figures S1-S6 for "Identification and Characterization of Fbxl22, a novel skeletal muscle atrophy-promoting E3 ubiquitin ligase"

**Supplemental Figure Legends**

**Supplement figure 1. Overexpression of Fbxl22-236 isoform leads to significantly increased expression of the novel Fbxl22-193 splice variant.** Mouse TA muscle was transfected with either empty vector (EV) or the Fbxl22-236 construct for 14 days. mRNA expression (fold change relative to EV) of **(A)** Fbxl22-236 and **(B)** Fbxl22-193 mRNA after 14 days of Fbxl22-236 overexpression. Statistical significance is depicted. Data presented as Mean ± SEM (N= 5/group).

**Supplemental figure 2. Biochemical analysis of protein degradation markers and sarcomeric proteins in Fbxl22-236 transfected muscles after 7 and 28 days.** (**A**) Representative immunoblot images for empty vector (EV) and Fbxl22-236 transfected muscles after 7 days (n=4/group). Total protein levels for **(B)** ubiquitin, **(C)** p62 and **(D)** LC3B II. Total protein levels for **(E)** dystrophin, **(F)** α-actinin (lower band), **(G)** desmin and **(H)** vimentin. (**I**) Representative immunoblot images for EV and Fbxl22-236 transfected muscles after 28 days (n=4/group). Total protein levels for **(J)** ubiquitin, **(K)** p62 and **(L)** LC3B II. Total protein levels for **(M)** dystrophin, **(N)** α-actinin (lower band), **(O)** desmin and **(P)** vimentin. Total protein loading was used as the normalization control for all blots. Statistical significance is depicted where present. Data presented as Mean ± SEM.

**Supplemental figure 3. Biochemical analysis of protein degradation markers and sarcomeric proteins in Fbxl22-193 transfected muscles after 7 and 28 days.** (**A**) Representative immunoblot images for empty vector (EV) and Fbxl22-193 transfected muscles after 7 days (n=4/group). Total protein levels for **(B)** ubiquitin, **(C)** p62 and **(D)** LC3B II. Total protein levels for **(E)** dystrophin, **(F)** α-actinin (lower band), **(G)** desmin and **(H)** vimentin. (**I**) Representative immunoblot images for EV and Fbxl22-193 transfected muscles after 28 days (n=4/group). Total protein levels for **(J)** ubiquitin, **(K)** p62 and **(L)** LC3B II. Total protein levels for **(M)** dystrophin, **(N)** α-actinin (lower band), **(O)** desmin and **(P)** vimentin. Total protein loading was used as the normalization control for all blots. Statistical significance is depicted where present. Data presented as Mean ± SEM.

**Supplemental figure 4. Biochemical analysis of protein degradation markers and sarcomeric proteins in lateral gastrocnemius muscles.** Lateral gastrocnemius muscles were transfected with either an empty vector (EV), Fbxl22-236 or Fbxl22-193 isoforms for 14 days. (**A**) Representative immunoblot images for EV and Fbxl22-236 transfected muscles (n=5/group). Total protein levels for **(B)** ubiquitin, **(C)** p62 and **(D)** LC3B II. Total protein levels for **(E)** dystrophin, **(F)** α-actinin (lower band), **(G)** desmin and **(H)** vimentin. (**I**) Representative immunoblot images for EV and Fbxl22-193 transfected muscles (n=5/group). Total protein levels for **(J)** ubiquitin, **(K)** p62 and **(L)** LC3B II. Total protein levels for **(M)** dystrophin, **(N)** α-actinin (lower band), **(O)** desmin and **(P)** vimentin. Total protein loading was used as the normalization control for all blots. Statistical significance is depicted where present. Data presented as Mean ± SEM.

**Supplemental figure 5. Biochemical analysis of protein degradation markers and sarcomeric proteins in medial gastrocnemius muscles.** Medial gastrocnemius muscles were transfected with either an empty vector (EV), Fbxl22-236 or Fbxl22-193 isoforms for 14 days. (**A**) Representative immunoblot images for EV and Fbxl22-236 transfected muscles (n=5/group). Total protein levels for **(B)** ubiquitin, **(C)** p62 and **(D)** LC3B II. Total protein levels for **(E)** dystrophin, **(F)** α-actinin (lower band), **(G)** desmin and **(H)** vimentin. **(I)** Representative immunoblot images for EV and Fbxl22-193 transfected muscles (n=5/group). Total protein levels for **(J)** ubiquitin, **(K)** p62 and **(L)** LC3B II. Total protein levels for **(M)** dystrophin, **(N)** α-actinin (lower band), **(O)** desmin and **(P)** vimentin. Total protein loading was used as the normalization control for all blots. Statistical significance is depicted where present. Data presented as Mean ± SEM.

**Supplemental figure 6. Knockdown of Fbxl22 expression after 3 days of denervation in wild type muscle.** Representative images for GFP fluorescence and laminin stain in Fbxl22 RNAi and empty vector transfected **(A)** lateral and **(B)** medial gastrocnemius muscles after 3 days of denervation (10x magnification; scale bar = 100µm). mRNA expression of Fbxl22-236 in **(C)** lateral and **(D)** medial gastrocnemius muscles transfected with empty vector or RNAi constructs. mRNA expression of Fbxl22-193 in **(E)** lateral and **(F)** medial gastrocnemius muscles transfected with empty vector or RNAi constructs. Data presented as Mean ± SEM (n=4-5/group). Φ depict P ≤ 0.05 verses empty vector group.

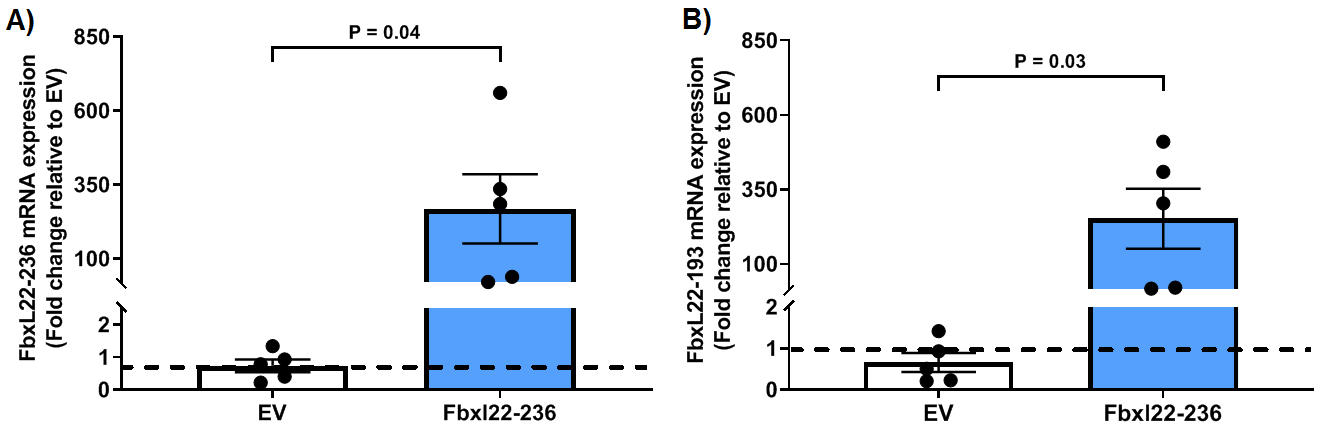

**Supplement figure 1.**

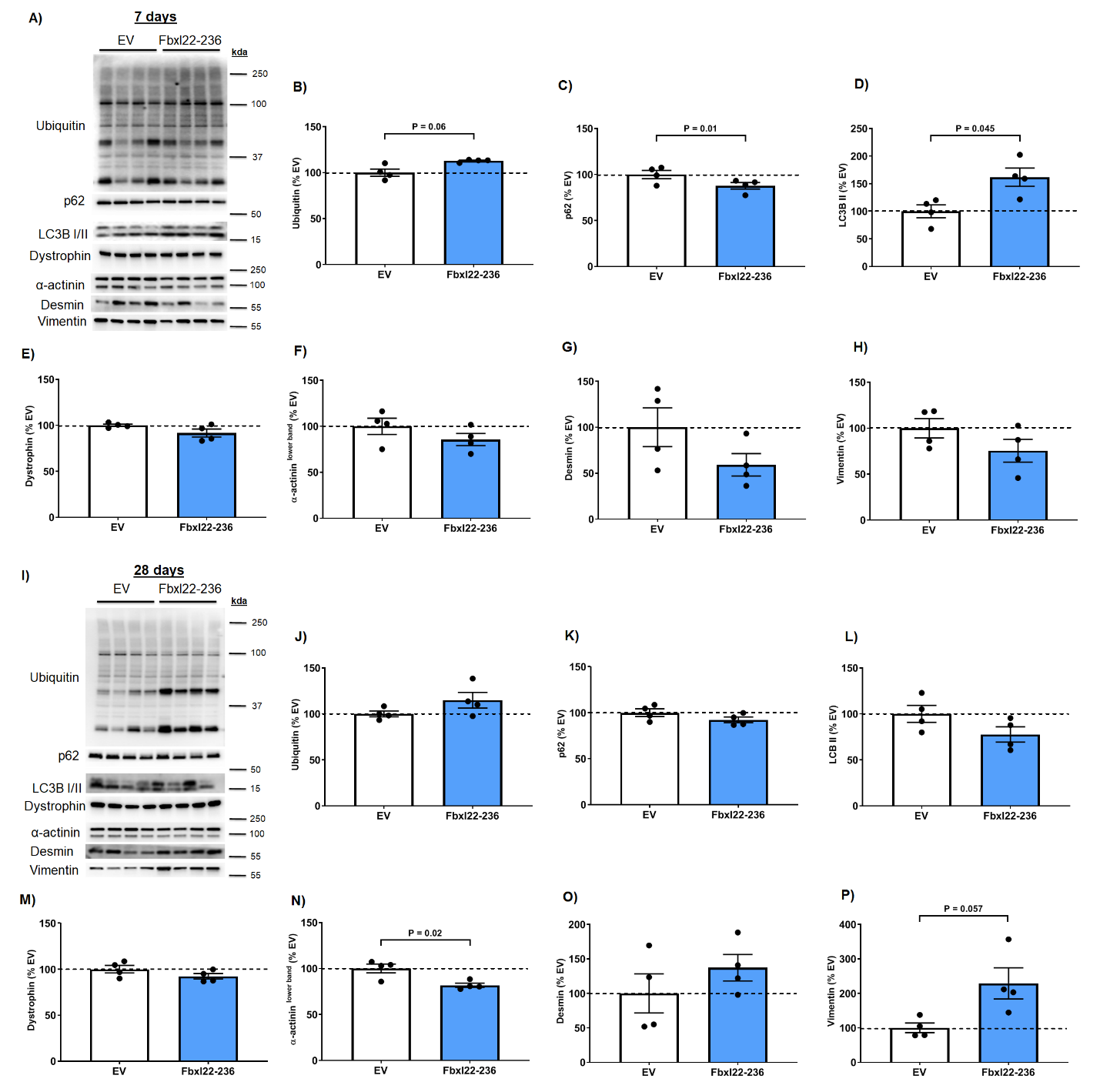

**Supplemental figure 2.**

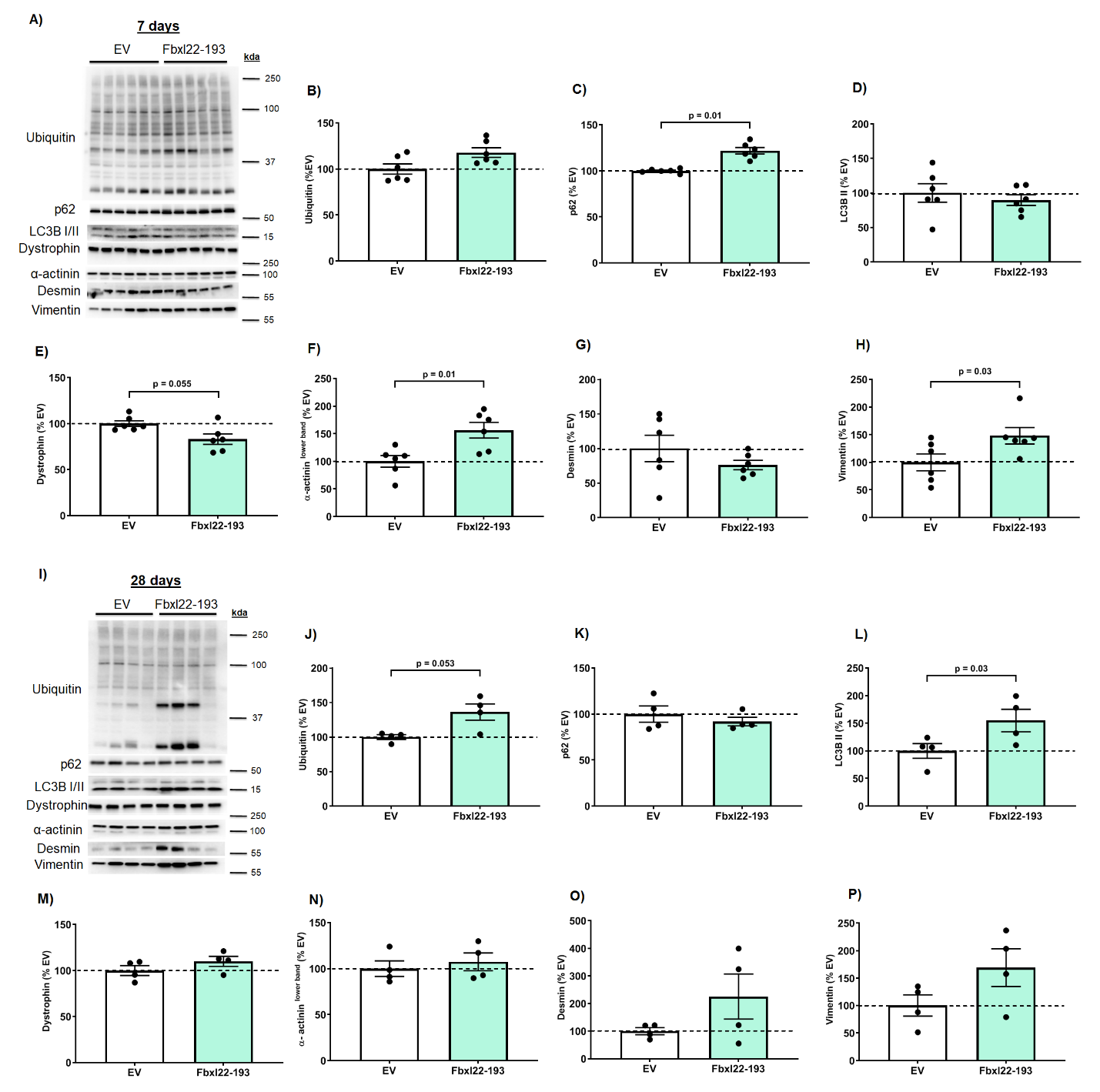

**Supplemental figure 3.**

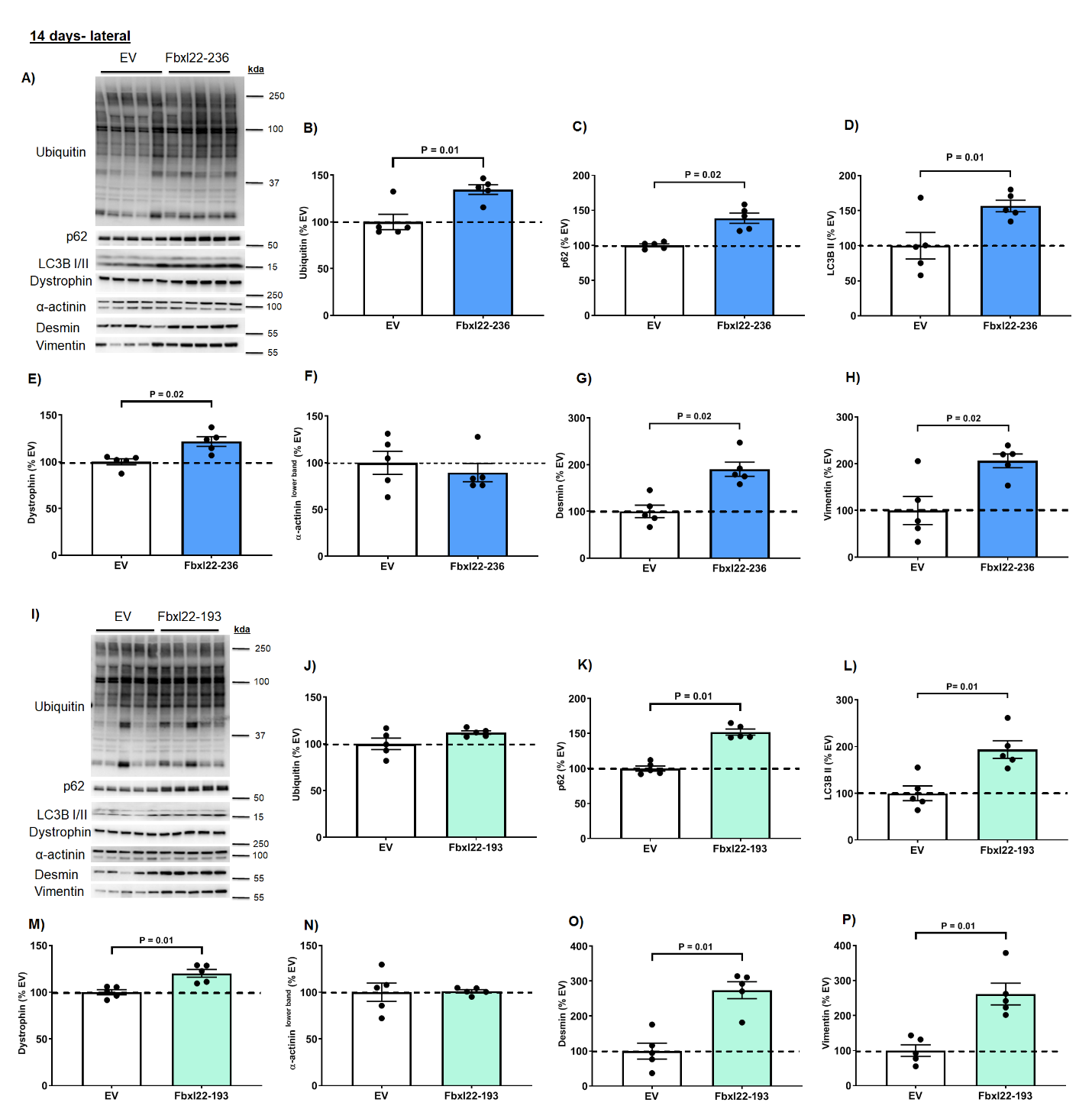

**Supplemental figure 4.**

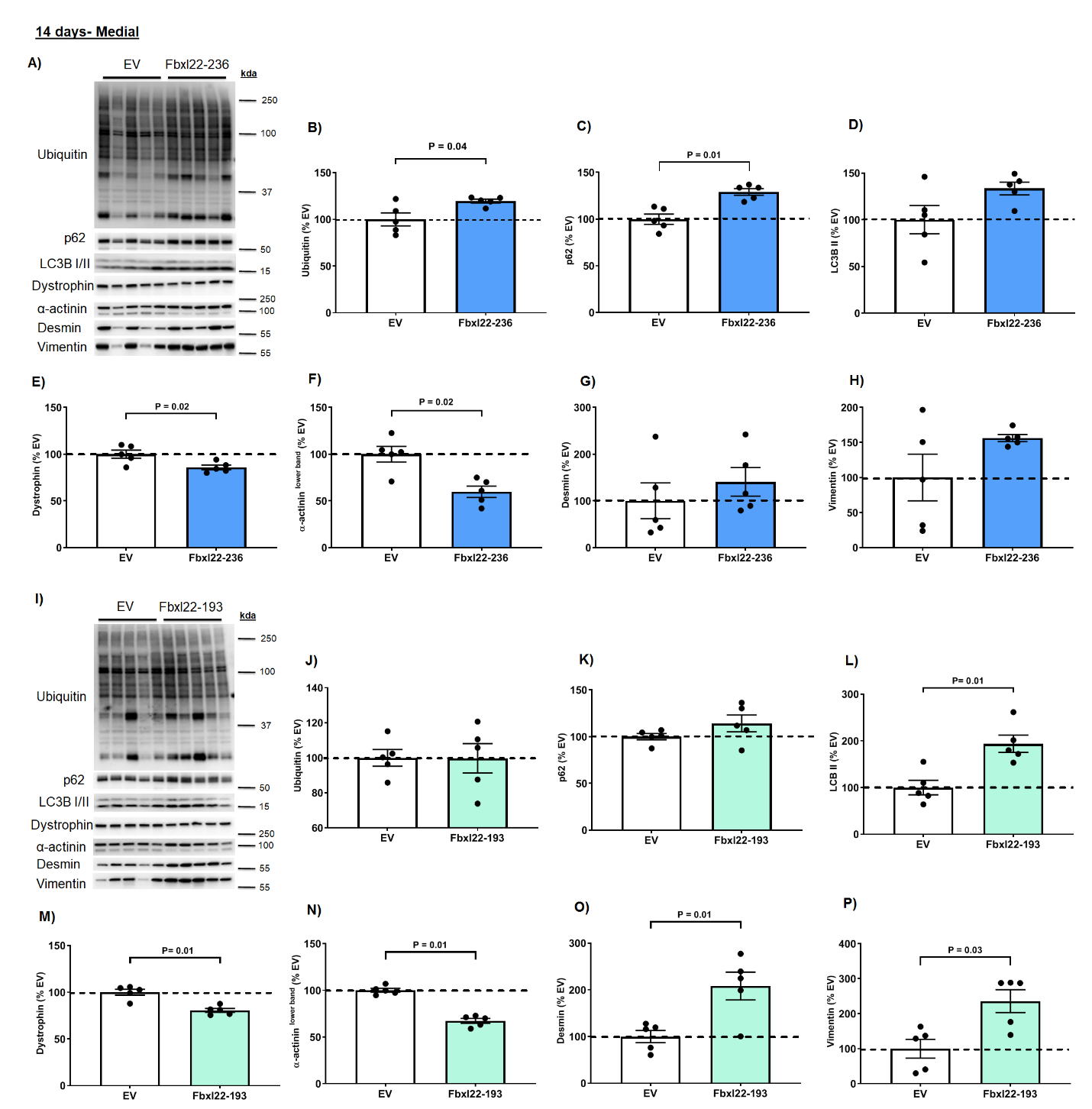

**Supplemental figure 5.**

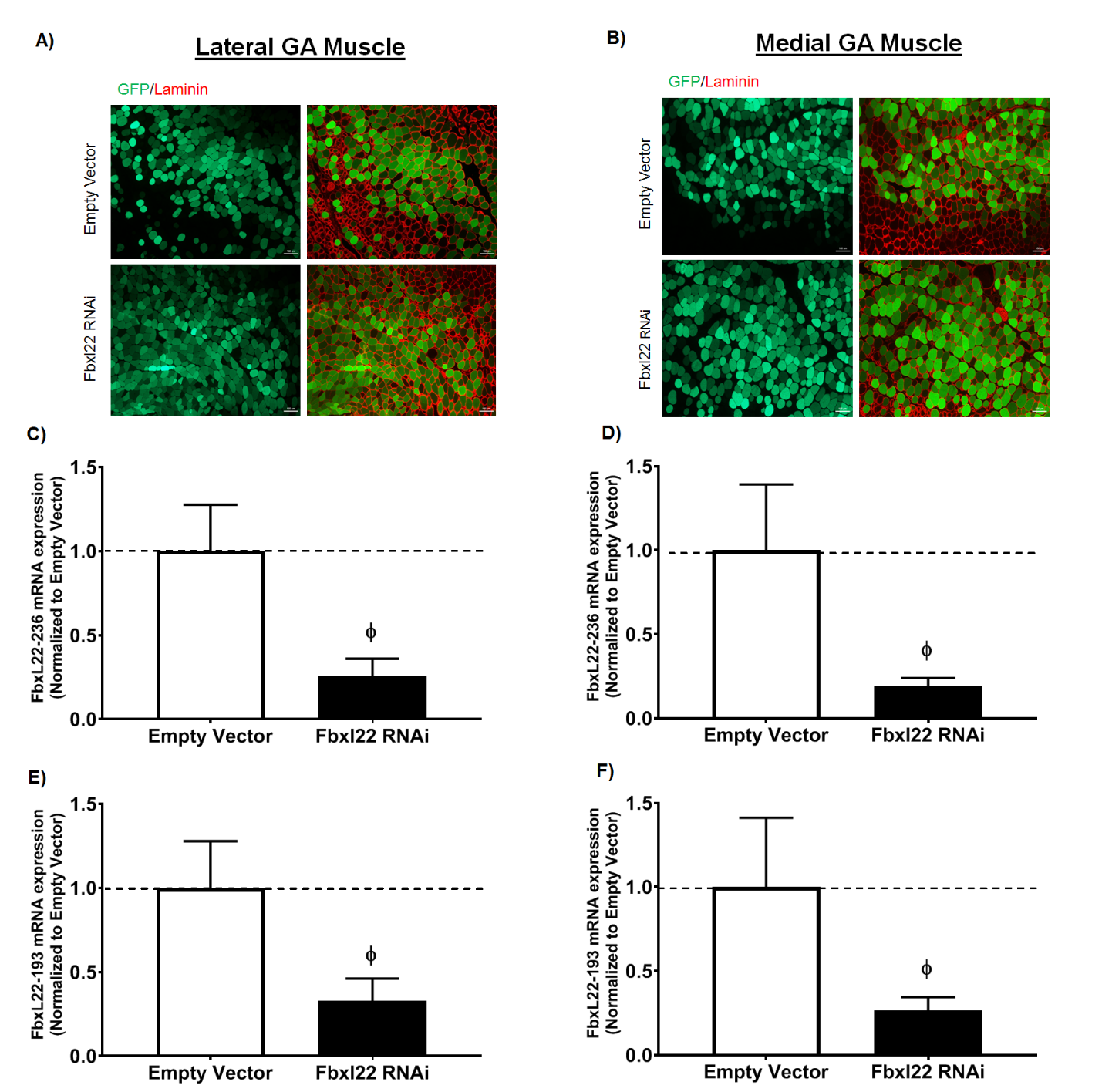

**Supplemental figure 6.**
